# Nucleolar Condensation Orchestrates rRNA-dependent Mobility and Spatiotemporally Enriches rDNA-binding of Human Chromatin Remodeler BRG1

**DOI:** 10.64898/2026.04.03.716269

**Authors:** Woei Shyuan Ng, Wilfried Engl, Ziqing Winston Zhao

## Abstract

As key regulators of genome access via nucleosome translocation/ejection, SWI/SNF chromatin remodeling complexes share a core ATPase/translocase subunit, BRG1, central to their activity. Despite recent discovery of spatially clustered intranuclear “hotspots” where various SWI/SNF remodelers preferentially bind, the mechanistic driving force underlying such heterogeneous organization remains unclear. Herein, we show that human BRG1 undergoes condensation *in vitro* and in live cell nucleus, mediated by its IDR-rich C-terminus (BRG1_C_). Intranuclear condensates of BRG1_C_ form across a wide range of (including endogenous) expression levels, are highly dynamic, and selectively partition into the fibrillar center of nucleolus, with their formation, localization and liquid-like properties governed primarily by patterned charge blocks in its sequence. Importantly, correlative single-molecule tracking and condensates mapping reveal rRNA-modulated constrained mobility and spatiotemporally enriched chromatin-binding to rDNA for BRG1_C_ specifically within nucleolar condensates. These findings unveil a condensation-mediated coupling between remodeler dynamics and nucleolar architecture, pointing to a potentially generic mechanism for organizing remodeling activity in both space and time.

## INTRODUCTION

In eukaryotic cells, the packaging of DNA into nucleosomes poses a topological challenge when the underlying target DNA needs to be accessed for diverse downstream processes. Such genome access is provided primarily through the actions of ATP-dependent chromatin remodelers (*1*). By facilitating nucleosome translocation/ejection, histone eviction or exchange, remodelers dynamically modulate chromatin accessibility, while acting within a larger regulatory network that synergizes with RNA polymerases, chromatin modifications and transcriptional activators/repressors to collectively regulate gene expression (*2*). Among the different subfamilies of chromatin remodelers, the SWI/SNF complex is one of the best characterized, of which three major subtypes (canonical BAF (cBAF), polybromo-associated BAF (PBAF), and non-canonical BAF (ncBAF)) are each assembled from 10-15 distinct subunits into a ∼1–1.5 MDa complex (*3*, *4*). However, common to all subtypes of the SWI/SNF complex is the core ATPase/translocase subunit, Brahma-related gene-1 (BRG1), which directly interacts with the nucleosome to effect the key translocation step of the chromatin remodeling process in an ATP-dependent manner (*5*, *6*). In addition to this central activity, BRG1 also associates with various other binding partners, enabling its incorporation into a diverse array of multi-subunit complexes that regulate transcriptional activation/repression, DNA replication and repair, and intracellular signalling pathways (*2*).

While the evolutionarily conserved ATPase/helicase domain of BRG1 has been extensively studied (*7*–*10*), a substantial portion (61.3%) of the protein sequence consists of intrinsically disordered regions (IDRs), particularly at its N- and C-termini (**Fig. 1A** and **Fig. S1**). As key mediators of weak multivalent interactions that facilitate biomolecular phase separation or condensation, IDRs are known to be critically involved in diverse intracellular processes by promoting the formation of localized microenvironments where biomolecules can be selectively concentrated for specialized activities (*11*, *12*). Moreover, such enrichment of intrinsic disorder in protein sequence is a common feature of many of the key SWI/SNF subunits (*13*), with IDR content up to 77.6% (**Fig. S1**), thereby pointing to their potentially varied involvements in the overall functionalities of the remodeling complexes. For example, the N-terminal IDR of the ARID1A subunit has been shown to play an important role in modulating cBAF targeting and activity through unique sequence features (*14*). Multiple SWI/SNF subunits can also be recruited by oncogenic FET fusion proteins into transcriptional co-condensates via heterotypic interactions (*15*, *16*). However, aside from these sporadic studies, the generic functional contribution of IDRs to SWI/SNF remodelers actions, particularly in relation to how they dynamically interact with chromatin to effect remodeling activity, remains unclear.

**Figure 1.**
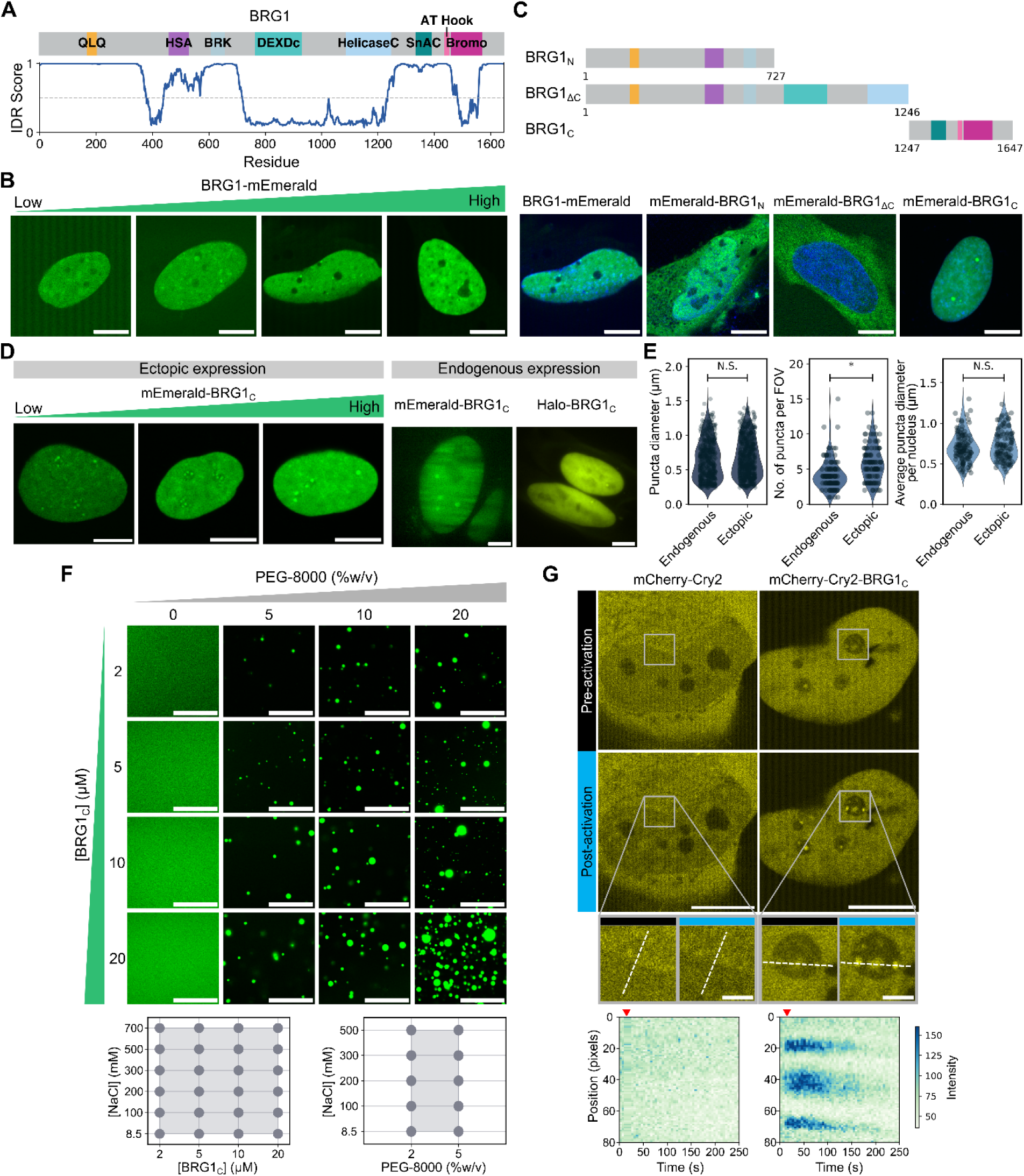
BRG1 forms condensates *in vitro* and in live human cell nucleus, mediated by its IDR-rich C-terminus. **(A)** Schematic of the domain organization of BRG1 and the IDR score predicted by AIUPRED across its sequence. **(B)** Confocal images of BRG1-mEmerald condensates formed in live Hela cells expressing BRG1 across different levels. **(C)** Schematics of various mEmerald-tagged BRG1 truncation variants generated (top) as well as confocal images showing their intracellular distributions in live Hela cells (bottom), indicating that the C-terminus is critical for its intranuclear localization. Cell nucleus is stained with Hoechst. **(D)** BRG1_C_ forms condensates across different levels of ectopic expression (left), as well as in CRISPR knock-in cell lines stably expressing mEmerald- or Halo-tagged BRG1_C_ at endogenous levels (right). **(E)** Quantification of BRG1_C_ puncta size, number of puncta per field of view (FOV), and mean puncta size per cell nucleus for both ectopically and endogenously expressing cells; each dot denotes either a single punctus (left) or a single cell (middle and right). *n* = 467 puncta from 111 cells (endogenous) and 587 puncta from 100 cells (ectopic). **(F)** *In vitro* droplet assay on purified mEmerald-10xHis-BRG1_C_ shows BRG1_C_ can form condensates across a wide range of protein, salt and PEG-8000 concentrations; pH was maintained at 8.0. **(G)** Confocal images of live Hela cells expressing either mCherry-Cry2 or mCherry-Cry2-BRG1_C_ before and after 7.2 s of photoactivation by 488-nm light (top); insets show zoomed-in intranuclear regions delineated in white boxes. Kymographs (bottom) show intensity changes across the white dotted lines in insets upon photoactivation (denoted by red arrowhead). Scale bar: 50 µm (**F**); 2 µm (**G** insets); 10 µm (all others).

Recently, the intranuclear diffusion and chromatin-binding dynamics of human SWI/SNF remodelers (including that of BRG1) have been systematically quantified using live-cell single-molecule imaging (*17*). Specifically, a multi-modal binding landscape in which preferential and repeated chromatin-binding of remodelers within heterogeneously distributed nanoscale “hotspots” across the nucleoplasm has been found to potentially drive sustained productive remodeling. However, the underlying mechanism for such spatio-temporal organization remains unknown. Hence, we hypothesize that IDR-mediated condensation of the common BRG1 subunit could serve as a mechanistic driving force for the heterogeneous intranuclear organization of SWI/SNF remodelers dynamics to selectively modulate remodeling activity. Herein, we combine quantitative live-cell imaging, single-molecule tracking, super-resolution mapping, *in vitro* biochemistry and bioinformatic analysis to show that human BRG1 undergoes condensation both *in vitro* and in live cell nucleus, mediated by its IDR-rich C-terminus. Importantly, these condensates preferentially partition into the fibrillar center phase of the nucleolus, in which BRG1 exhibits constrained mobility modulated by rRNAs as well as spatiotemporally enriched chromatin-binding to rDNA at the single-molecule level. Collectively, these findings unveil a unique condensation-mediated structural association as well as dynamic coupling between BRG1 and the nucleolus, which could potentially serve as a generic mechanism for organizing remodeling activity in both space and time.

## RESULTS

### BRG1 forms condensates *in vitro* and in live human cell nucleus, mediated by its IDR-rich C-terminus

In light of the high level of intrinsic disorder in BRG1 as predicted by both AIUPRED and AlphaFold DB (**Fig. 1A** and **Fig. S1**), we first examined if it can undergo phase separation in live human cells. Indeed, HeLa cells ectopically expressing mEmerald-tagged BRG1 form droplet-like puncta characteristic of liquid-state condensates inside the cell nucleus across a wide range of expression levels (**Fig. 1B**), with the propensity for condensates formation positively correlating with intranuclear BRG1 level (**Fig. S2**). In light of the presence of two major IDRs in the N- and C-termini of BRG1, we generated three different mEmerald-tagged truncation variants of BRG1 for N-terminus only (BRG1_N_), C-terminus only (BRG1_C_), or with the C-terminus truncated (BRG1_ΔC_) (**Fig. 1C**). Among them, only BRG1_C_ showed similar nuclear localization and condensates formation as the full-length BRG1; in contrast, BRG1_N_ and BRG1_ΔC_ could not be specifically localized to the nucleus, with BRG1_N_ showing a diffuse distribution across both the nucleus and cytoplasm while BRG1_ΔC_ was largely excluded from the nucleus (**Fig. 1C**). Moreover, similar intranuclear condensates formation was observed both in cells ectopically expressing mEmerald-BRG1_C_ across different levels and in CRISPR knock-in cell lines stably expressing either mEmerald- or Halo-tagged BRG1_C_ at endogenous levels (**Fig. 1D**), which exhibited comparable puncta size and number per nucleus (**Fig. 1E**). The endogenous BRG1_C_ level was also found to fall within the range of ectopic expression levels observed (**Fig. S3**), hence ruling out the possibility of expression-level-associated artifacts. These findings collectively show that the intranuclear condensation behavior observed for BRG1_C_ can largely recapitulate that of full-length BRG1.

As a further validation for our live-cell observations, *in vitro* droplet assay was performed on biochemically purified BRG1_C_ protein, which forms droplets *in vitro* under physiologically relevant pH, ionic strengths, as well as the presence of molecular crowding agent mimicking the dense intranuclear environment (**Fig. 1F**). Similar to the *in cellulo* case, the *in vitro* condensation propensity also positively correlates with BRG1_C_ concentration. Moreover, HeLa cells expressing mCherry-tagged Cry2-BRG1_C_, adapted from the “optoDroplets” system capable of light-activated control of intracellular phase transitions (*18*), showed rapid clustering of BRG1_C_ into newly formed puncta as well as enhanced recruitment of BRG1_C_ into existing puncta upon brief blue-light photoactivation (**Fig. 1G**). These, together with the fact that BRG1_C_ condensates formation is independent of the fluorescent protein or tag to which it is fused (**Fig. S4**), indicate that the intranuclear localization and condensation of BRG1 are both mediated by its IDR-rich C-terminus.

### BRG1_C_ condensates formation, localization and liquid-like properties are governed by patterned charge blocks

To quantitatively characterize the dynamic and liquid-like properties of intranuclear BRG1_C_ condensates, fluorescence recovery after photobleaching (FRAP) measurement was first performed in live Hela cells. BRG1_C_ condensates exhibited a short recovery half-time of τ_1/2_ ≈ 0.4 s and a recovery fraction of ∼0.95 (**Fig. 2A**), indicating that they are highly dynamic and undergo rapid molecular exchange with the surrounding intranuclear milieu. Upon contact, BRG1_C_ droplets can also undergo fusion over a timescale of minutes (**Fig. 2B**), another hallmark of liquid-like condensates. However, intranuclear BRG1_C_ condensates were largely recalcitrant to 1,6-hexanediol (1,6-HD) treatment across a range of concentrations for up to 10 minutes (**Fig. 2C** and **Fig. S5**), suggesting that the weak hydrophobic interactions disrupted by 1,6-HD are not the main driving force for BRG1_C_ condensation.

**Figure 2.**
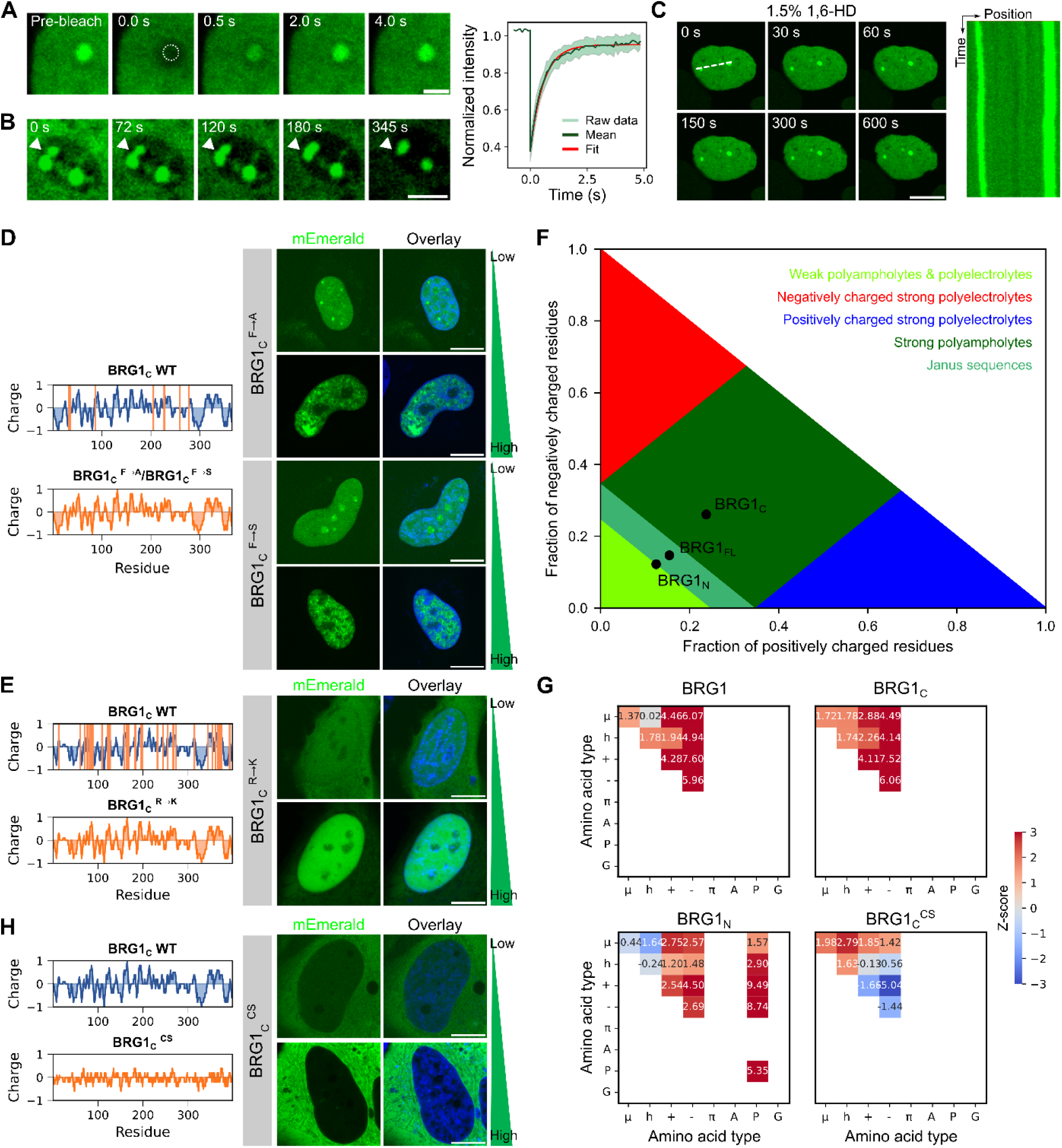
BRG1_C_ condensates formation, localization and liquid-like properties are governed primarily by patterned charge blocks. **(A)** A representative FRAP measurement on mEmerald-BRG1_C_ condensate (circled in white dotted line) in a live Hela cell (left), which exhibits a fast recovery half-time of ∼0.4 s and a recovery fraction of ∼0.95 (right), both indicative of highly dynamic structures. **(B)** Confocal images of mEmerald-BRG1_C_ condensate droplets in a live Hela cell undergoing fusion (indicated by arrowhead) over a period of 345 s upon treatment with 2.0% 1,6-hexanediol (1,6-HD). **(C)** Confocal images of mEmerald-BRG1_C_ condensates in a live Hela cell that persist for up to 10 minutes upon treatment with 1.5% 1,6-HD. Kymograph on the right shows fluorescence intensity change across the white dotted line. **(D)** Das-Pappu phase diagram of states generated with CIDER allocates full-length BRG1 (BRG1_FL_), BRG1_N_ and BRG1_C_ to regions corresponding to Janus sequences, weak polyampholytes and strong polyampholytes, respectively. **(E)** Z-score matrices of sequence patterning parameters calculated by NARDINI for BRG1, BRG1_N_ and BRG1_C_ (both wild-type and charge-scrambled mutant). Residues are grouped according to polar or μ (S, T, N, Q, C, H), hydrophobic or h (I, L, M, V), positively charged or + (R, K), negatively charged or − (E, D), aromatic or π (F, W, Y), alanine (A), proline (P), and glycine (G). **(F** to **H)** Plots of linear net charge per residue of wild-type BRG1_C_ and BRG1_C_ ^F→A^ / BRG1_C_ ^F→S^ (**F)**, BRG1_C_ ^R→K^ (**G**) and BRG1_C_ ^CS^ (**H**) mutants, as well as corresponding confocal images of HeLa cells expressing each mutant (tagged with mEmerald). Cell nucleus is stained with Hoechst. Orange vertical lines in the wild-type BRG1_C_ plot denote positions with mutated residues for each mutant. Scale bar: 2 µm (**A** and **B**); 10 µm (all others).

In order to elucidate the specific “molecular grammar” that governs BRG1_C_ condensation, we first examined the potential roles of two key “stickers” in the “sticker-and-spacer” framework for mediating biomolecular condensation (other than hydrophobic interactions): π−π interactions mediated by aromatic residues (*e.g.* phenylalanine), and cation−π interactions (a key mediator of which being arginine) (*19*, *20*). Given that the BRG1_C_ sequence contains a total of 7 phenylalanine residues and 37 arginine residues, we generated three BRG1_C_ mutants by mutating either all phenylalanine residues to alanine or serine (F→A or F→S mutant), or all arginine residues to lysine (R→K mutant) (**Fig. 2D** and **E**). While cells expressing low levels of the BRG1_C_^F→A^ or BRG1_C_^F→S^ mutant could still form condensates, they formed large aggregates across the nucleoplasm at high expression levels (**Fig. 2D**), pointing to the role of aromatic residues in modulating the fluid properties of BRG1_C_ condensates to prevent aberrant aggregation. On the other hand, the BRG1_C_^R→K^ mutant, in which the arginine-mediated cation−π interactions are weakened, exhibited diminished ability to specifically localize to the nucleus at low expression levels; while such intranuclear targeting ability is partially recovered at high expression levels, condensates formation of the mutant nonetheless becomes significantly reduced (**Fig. 2E**).

Moreover, CIDER and NARDINI, two commonly used computational algorithms for analyzing the physico-chemical properties of and sequence patterns in IDRs (*21*, *22*), both identified a strong patterning of opposite charge blocks in the sequence of BRG1_C_ characteristic of strong polyampholytes (**Fig. 2F** and **G** and **Fig. S6**). In light of that, we further generated a charge-scrambled (CS) mutant that scrambles the opposite charge blocks in BRG1_C_ without altering its net charge, and found that the BRG1_C_^CS^ mutant completely abolishes both its intranuclear localization and condensate formation (**Fig. 2H** and **Fig. S7**). Therefore, electrostatic interactions, particularly in the form of a blocky pattern of opposite charges characteristic of polyampholytes, serve as the main driving force for BRG1_C_ condensation.

### BRG1_C_ condensates selectively partition into the fibrillar center phase of nucleolus and can be modulated by rRNAs

Importantly, intranuclear BRG1_C_ condensates were observed to preferentially localize to the nucleolus, which is known to be organized as a multi-layered condensate consisting of three distinct phases (from the outside to the inside): granular component (GC), dense fibrillar component (DFC), and fibrillar center (FC) (*23*). To pinpoint the specific spatial relationship between BRG1_C_ condensates and these nucleolar phases, Halo-BRG1_C_ was co-expressed in live HeLa cells with each of the following key nucleolar markers: NPM1 (for GC), fibrillarin (for DFC), and RPA194 (a key catalytic subunit of RNA polymerase I (Pol I)), UBF and RPA43 (for FC), each tagged with GFP (**Fig. 3A**). Colocalization analysis indicated that BRG1_C_ condensates specifically partition into the FC phase of the nucleolus, nested within the DFC and GC phases. Similar colocalization patterns were also observed with dual-color immunofluorescence imaging (**Fig. S8**). Such selective partitioning strongly suggests some form of functional coupling between BRG1 condensates and the nucleolar activities within the FC phase.

**Figure 3.**
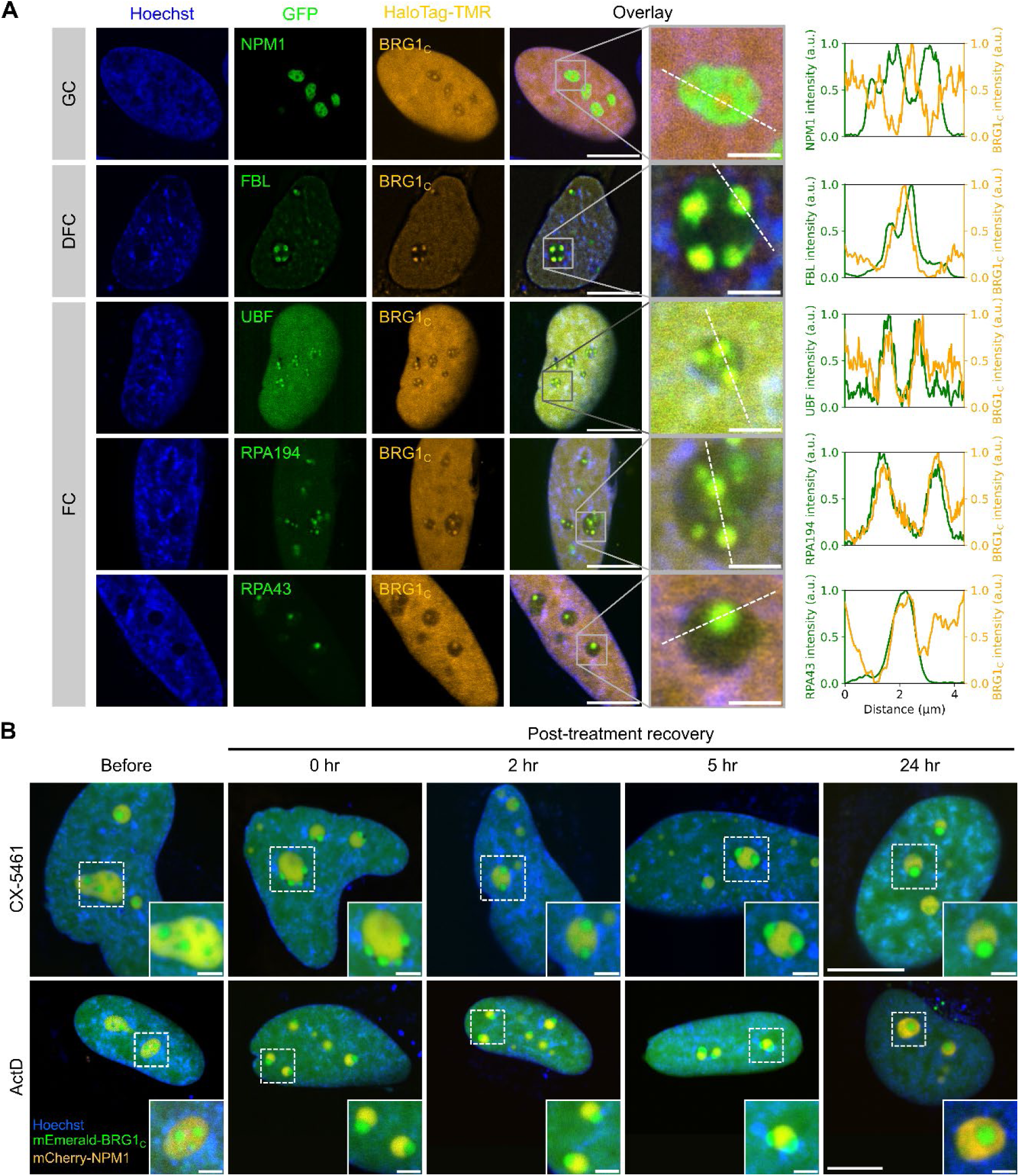
BRG1_C_ puncta colocalize with the fibrillar center phase of nucleolus, and can be modulated by rRNAs. **(A)** Confocal images of live Hela cells co-expressing Halo-BRG1_C_ (labelled with TMR dye) with each of the key markers for the three nucleolar phases (labelled with GFP): NPM1 of granular component (GC), fibrillarin (FBL) of dense fibrillar component (DFC), and UBF, RPA194 and RPA43 of fibrillar center (FC). Cell nucleus was stained with Hoechst. Insets show zoomed-in regions of the overlay images delineated in white boxes, with the fluorescence intensity changes across the white dotted lines plotted on the right. **(B)** Confocal images of live HeLa cells co-expressing mEmerald-BRG1_C_ and mCherry-NPM1 before and after treatment with either CX-5461 (top row) or actinomycin D (ActD) (bottom row). Cell nucleus was stained with Hoechst. Insets show zoomed-in view of the ultrastructural changes undergone by BRG1_C_ condensates in relation to nucleolus, delineated in white boxes. Scale bar: 10 µm (all main images); 2 µm (all insets).

Moreover, in light of the fact that the nucleolus is the primary site of ribosome biogenesis where ribosomal DNA (rDNA) is located and transcribed at the FC/DFC boundary (*24*–*26*), we treated the cells with either CX-5461 (a direct and selective inhibitor of ribosomal RNA (rRNA) transcription via inhibiting Pol I inside the nucleolus (*27*)) or actinomycin D (ActD, which generically inhibits transcription mediated by all RNA polymerases (*28*)). Importantly, both treatments caused BRG1_C_ condensates to form “nucleolar cap”-like structures (**Fig. 3B**), a well-known architectural re-organization associated with nucleolar stress in which the interior FC phase flips out and relocates to the nucleolar periphery (*23*). Importantly, the normal positioning of BRG1_C_ condensates within the nucleolar interior can be recovered within 24 hours from ActD treatment, whereas the effect of CX-5461 treatment is irreversible during this timeframe (**Fig. 3B**). All of these collectively indicate that Pol I-mediated rRNA transcription within the nucleolus plays a crucial role in modulating both the architectural organization as well as the liquid-like properties of BRG1_C_ condensates, and points to the potential functional implications of such preferential remodeler–nucleolus association for BRG1’s dynamic interactions with rDNA at the FC/DFC boundary.

### BRG1_C_ within the nucleolar condensates exhibits more constrained mobility

In order to quantitatively interrogate how nucleolar condensation impacts BRG1’s intranuclear mobility and chromatin-binding dynamics, we performed live-cell single-molecule tracking (SMT) under HILO illumination on Halo-BRG1_C_ labeled with JF_549_ dye, while simultaneously monitoring the intranuclear DNA background stained with Hoechst (**Fig. 4A**). We first performed fast tracking (with 5.5 ms frame time) to probe the more transient diffusional dynamics of BRG1_C_ (**Fig. 4B** and **Movie S1**), and analyzed the SMT trajectories with the “Spot-On” framework (*29*). The displacement histogram obtained can only be satisfactorily fitted with a three-state model (**Fig. 4C**), with each state having a distinct diffusion coefficient *D* (**Fig. 4D**), hence indicating that a minimum of three mobility modes is required to fully describe the intranuclear dynamics of BRG1_C_. The fast-diffusion and slow-diffusion modes, with mean diffusion coefficients of 0.82 µm^2^s^-1^ and 5.2 µm^2^s^-1^ respectively, likely correspond to the diffusion of individual BRG1_C_ subunit and of the assembled remodeler complex, while the slowest mode (with a mean diffusion coefficient of 0.13 µm^2^s^-1^) likely corresponds to the chromatin-bound remodeler complex. Importantly, all three modes were previously observed for the full-length BRG1 (*17*) (**Fig. 4D**), hence suggesting that the full spectrum of BRG1’s intranuclear mobility is retained in BRG1_C_.

**Figure 4.**
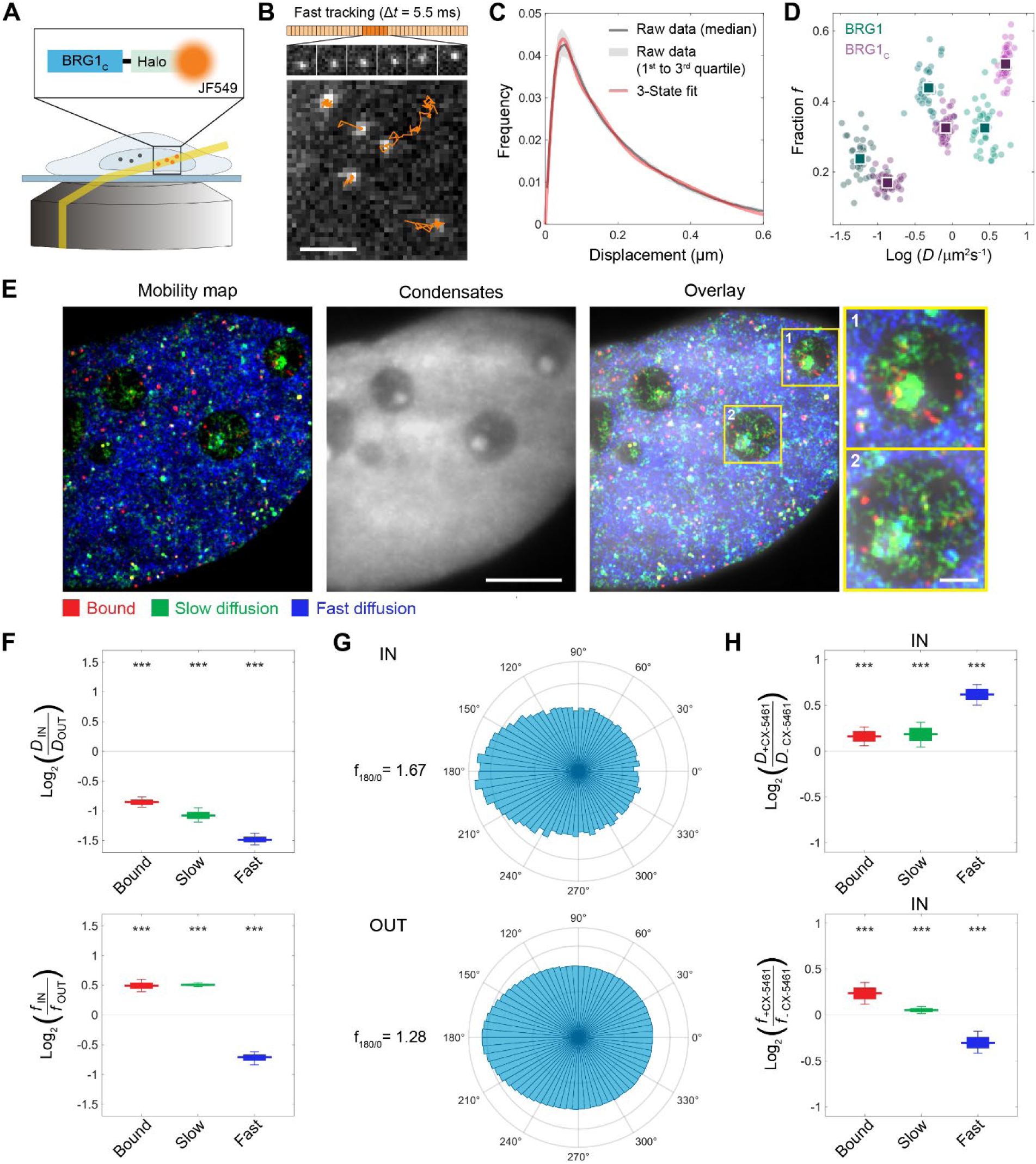
Correlative single-molecule tracking and condensates mapping show that BRG1_C_ undergoes more constrained diffusion within nucleolar condensates. **(A)** Schematic of single-molecule tracking (SMT) of Halo-tagged BRG1_C_ dynamics in live Hela cells under HILO illumination. **(B)** A representative frame from a SMT movie superimposed with trajectories of individual diffusing Halo-BRG1_C_ molecules (orange) acquired under fast tracking. **(C)** A representative displacement histogram derived from fast-tracking trajectories (as in **(B)**) that can only be satisfactorily fitted with a three-state model. **(D)** Scatter plots of the diffusion coefficient (*D*) and the corresponding fraction (*f*) associated with each of the three mobility modes resolved from fast-tracking trajectories for full-length BRG1 (green) and BRG1_C_ (pink), overlaid on top of each other. **(E)** Super-resolved STAR map for each of the three mobility modes of BRG1_C_ (left, color-coded based on diffusion coefficient values) and the intranuclear distribution of BRG1_C_ condensates in the same cell (middle), as well as an overlay of the two (right). Insets show zoomed-in regions of the overlay images delineated in yellow boxes where the slow-diffusion and bound modes are significantly enriched. **(F)** Box-and-whisker plots of fold differences (on a log_2_ scale) between inside and outside the nucleolar condensates for diffusion coefficient (*D*, top) and corresponding fraction (*f*, bottom) associated with each of the three mobility modes of BRG1_C_. **(G)** Diffusional anisotropy analysis of fast-tracking trajectories from both inside (top) and outside (bottom) the nucleolar condensates indicates that BRG1_C_ within the condensates has a higher tendency to move backward between adjacent steps. **(H)** Box-and-whisker plots of fold changes (on a log_2_ scale) upon CX-5461 treatment in *D* (top) and *f* (bottom) associated with each of the three mobility modes of BRG1_C_ from trajectories inside the nucleolar condensates only. Scale bar: 2 µm **(B)**; 5 µm **(E)**. In **D**, each dot denotes a single cell, and squares denote mean values. *n* = 61 (BRG1_C_) and 39 (BRG1) cells for **D**, 7,090,454 trajectories from 31 cells for **F** and **G**, and 174,853 trajectories (–CX-5461) and 173,071 trajectories (+CX-5461) from 28 cells each for **H**. Data for full-length BRG1 in **D** adapted from (*17*).

Moreover, to furnish spatial contexts to such temporal dynamics, we previously devised a super-resolved strategy for mapping intranuclear mobility using SMT trajectories, termed STAR, capable of revealing the spatial distributions of various modes of remodeler dynamics (*e.g.* in the form of nanoscale “hotspots”) across the nucleoplasm (*17*). Performing STAR mapping on BRG1_C_ based on diffusion coefficient values obtained from the fast-tracking measurements, and correlating the resulting mobility map for each of the three modes with the intranuclear distribution of BRG1_C_ condensates (**Fig. 4E**), we observed a significant enrichment of the slow-diffusion and bound modes of BRG1_C_ within the nucleolus as compared to rest of the nucleoplasm. To quantify this trend more accurately, we isolated trajectories within the nucleolar condensates (denoted as ‘IN’) versus those outside the condensates (denoted as ‘OUT’), and found that BRG1_C_ within the nucleolar condensates exhibited significant reductions in the diffusion coefficients associated with all three mobility modes (by an average of 45–64%) (**Fig. 4F**), in line with the more viscous and crowded environment known to exist within the FC phase (*30*). In addition, the fraction associated with the fast-diffusion mode is reduced by an average of 39% within the nucleolar condensates, while that associated with the slow-diffusion and bound modes are increased by an average of 42% and 41%, respectively, pointing to a shift in the molecular partitioning of BRG1_C_ within the condensates towards these slower mobility modes. Moreover, diffusional anisotropy analysis on SMT trajectories found that BRG1_C_ within the nucleolar condensates exhibits a higher tendency to move backward between adjacent steps (as evidenced by a f_180/0_ ratio of 1.67) compared to those outside the condensates (for which f_180/0_ drops to 1.28) (**Fig. 4G**), indicating that BRG1_C_ within the condensates undergoes more constrained diffusion characteristic of highly crowded molecular environments. Finally, CX-5461 treatment led to higher diffusion coefficients for all three mobility modes of BRG1_C_ within the nucleolar condensates (**Fig. 4H**), suggesting that inhibiting rRNA transcription makes BRG1_C_ condensates more fluid, in line with the previous finding of a more liquid-like FC phase upon CX-5461-induced nucleolar cap formation (*31*). Such effect, however, can no longer be discerned if SMT trajectories from both inside and outside the nucleolar condensates are pooled together (**Fig. S9**). Collectively, these findings demonstrate that key dynamic properties of BRG1_C_ can be modulated in the condensates by rRNAs, and point to a potential “scaffolding” role provided by rRNAs for anchoring the remodeler within the FC phase of nucleolus, likely with functional consequences.

### Remodeler–nucleolus coupling spatiotemporally enriches chromatin-binding of BRG1_C_ to rDNA within the condensates

Finally, in order to probe the chromatin-binding dynamics of BRG1_C_ (which takes place on a longer timescale than diffusion), we further performed SMT under slow tracking (with 300 ms frame time), at which the rapidly diffusing BRG1_C_ molecules were mostly blurred out (**Fig. 5A** and **Movie S2**). Upon isolating binding events unambiguously from diffusion in the observed trajectories using a previously devised strategy (see **Materials and Methods** for details) (*17*), we found that the survival time distributions (*32*) of bound BRG1_C_ molecules were better fitted with a three-state model as opposed to a two-state model (**Fig. 5B**). Further validation with the genuine rate identification method (GRID) (**Fig. 5C**), an analytical framework capable of inferring the number, kinetics and amplitudes of superimposed processes from multi-component reaction systems in an unbiased manner (*33*), confirms that three distinct modes are needed to fully describe the binding dynamics of BRG1_C_: a transient binding mode with a mean residence time of 0.96 s, as well as two stable binding modes with mean residence times of 4.3 s and 18.6 s, respectively (**Fig. 5D**). Again, all three modes were previously observed for full-length BRG1 incorporated in the fully assembled SWI/SNF remodeler complex (*17*) with similar residence times (**Fig. 5D** and **Fig. S10**), thereby suggesting that BRG1’s chromatin-binding ability (within the overall SWI/SNF complex) is not compromised in BRG1_C_. Electrophoretic mobility shifty assay also confirmed that BRG1_C_ can bind to nucleosomal DNA, the presence of which can enhance BRG1_C_ condensation *in vitro* (**Fig. S11**). Moreover, treating the cells with SAHA, a histone deacetylase inhibitor known to enhance histone acetylation and decondense chromatin (*34*), led to quantitative alterations in the residence time, fraction and density of stable binding events (**Fig. S12**), in line with the previously reported role of the bromodomain within BRG1 C-terminus in contributing to binding hyperacetylated and decondensed chromatin (*35*). Together, these results ascertain that the intranuclear binding dynamics we observed for BRG1_C_ (including those inside the condensates) indeed corresponds to binding to chromatin DNA (and within the nucleolus, to rDNA).

**Figure 5.**
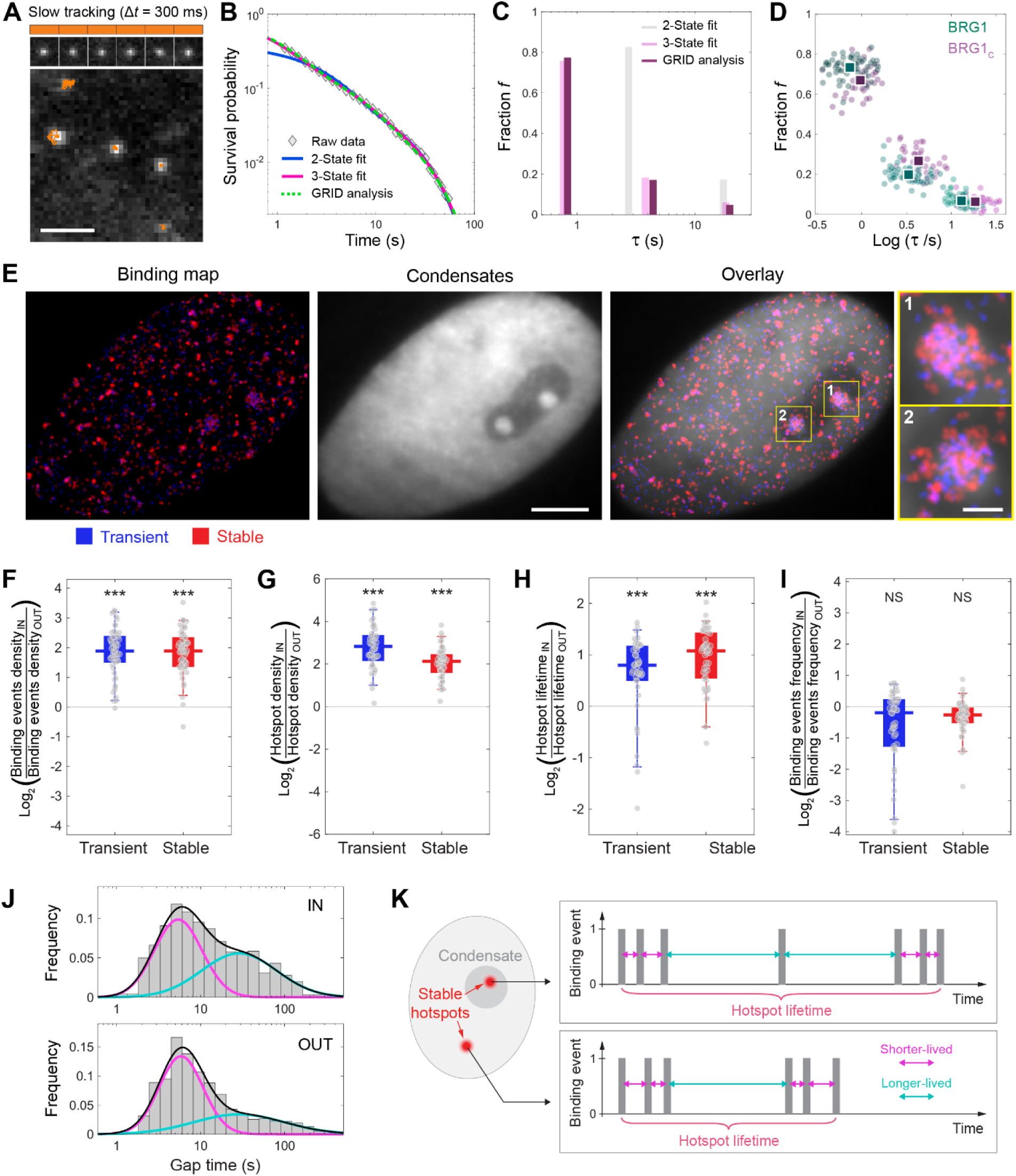
Nucleolar condensation enriches chromatin-binding of single BRG1_C_ molecules to rDNA in both space and time. **(A)** A representative frame from a SMT movie superimposed with trajectories of individual bound Halo-BRG1_C_ molecules (orange) acquired under slow tracking. **(B)** A representative survival probability distribution derived from slow-tracking trajectories (as in **(A)**), fitted with either two-state model, three-state model, or GRID analysis. **(C)** GRID analysis of slow-tracking trajectories yielded three distinct chromatin-binding modes, with residence times in much better agreement with those obtained from three-state model than those from two-state model. **(D)** Scatter plots of the residence time (*τ*) and the corresponding fraction (*f*) associated with each of the three chromatin-binding modes resolved from slow-tracking trajectories for full-length BRG1 (green) and BRG1_C_ (pink), overlaid on top of each other. **(E)** Super-resolved STAR map for transient and stable (combined) modes of BRG1_C_ chromatin-binding (left, color-coded based on residence times) and the intranuclear distribution of BRG1_C_ condensates in the same cell (middle), as well as an overlay of the two (right). Insets show zoomed-in regions of the overlay images delineated in yellow boxes where both binding modes are significantly enriched. **(F** to **I)** Box-and-whisker plots of fold differences (on a log_2_ scale) between inside and outside the nucleolar condensates for binding events density **(F)**, hotspots density **(G)**, binding events frequency **(H)**, and hotspots lifetime **(I)** for both transient and stable (combined) chromatin-binding of BRG1_C_. **(J)** Histogram of gap time between adjacent stable binding events of BRG1_C_ within a hotspot either inside (top) or outside (bottom) the nucleolar condensates, fitted with a double-Gaussian fit (black, with the shorter-lived component in pink and the longer-lived component in green). **(K)** Schematic illustrating how the temporal structure of stable binding hotspots of BRG1_C_ is modulated within the nucleolar condensates. Scale bar: 2 µm **(A)**; 5 µm **(E)**. In **D**, each dot denotes a single cell, and squares denote mean values. *n* = 58 (BRG1_C_) and 40 (BRG1) cells for **D**, 53 cells for **F** to **I**, and 11,660 gap times from 53 cells for **J**. Data for full-length BRG1 in **D** adapted from (*17*).

We next mapped the spatial distribution of BRG1_C_’s chromatin-binding based on residence times obtained from the slow-tracking measurements, and again correlated the binding map with its intranuclear condensates. Similar to full-length BRG1 (*17*), the chromatin-binding events of BRG1_C_ for both the transient mode and the stable mode (combining the two sub-modes) cluster into heterogeneously distributed binding “hotspots” across the nucleoplasm (**Fig. 5E**), where BRG1_C_ binds preferentially and repeatedly. In particular, both binding modes are spatially enriched within the nucleolar condensates, as manifested by the densities (per unit area) of binding events (enriched by an average of 3.7 folds) as well as of binding hotspots (enriched by an average of 4.4–7.1 folds) (**Fig. 5F** and **G**). In the temporal domain, the lifetimes of hotspots (defined as the duration spanned by all binding events within a hotspot) for both transient and stable binding are also significantly prolonged inside the condensates by an average of 1.7–2.1 folds (**Fig. 5H**), although the frequency of binding events per unit time remains largely unchanged between inside and outside the condensates for both modes (**Fig. 5I**). In order to better understand the temporal organization of binding events within the hotspots, we further quantified the gap time between adjacent binding events for both modes, and found that the gap time between stable binding events both inside and outside the condensates exhibits a bimodal distribution with two temporally distinct peaks at 5.4–5.8 s and 28.7–28.8 s, respectively; however, the relative frequency of the longer-lived gap time is substantially higher inside the condensates as compared to outside (**Fig. 5J**). In contrast, the gap time between transient binding events exhibits a more homogeneous distribution without such clear temporal separation (**Fig. S13**). As such, while key parameters of the temporal structure of stable binding hotspots remain largely unaffected by condensation, the increased frequency of the longer-lived gap time within the condensates allows more disparate stable binding events to be temporally adjoined into a hotspot with prolonged lifetime (**Fig. 5K**). In these two ways, nucleolar condensation enriches stable chromatin-binding of BRG1_C_ in both space and time, potentially serving as a strategy to more effectively carry out remodeling activity on rDNA within the nucleolus.

## DISCUSSION

In this work, we undertook a multi-pronged approach to quantitatively dissect the intranuclear condensation of BRG1, the core ATPase/translocase of SWI/SNF remodeler complexes, as well as the interplay between its chromatin-binding dynamics and nucleolar organization in live human cells. We found that the intranuclear localization and condensation of BRG1 are mediated by its IDR-rich C-terminus (BRG1_C_), which can largely recapitulate the condensation behavior of full-length BRG1 across a wide range of (including endogenous) expression levels. Importantly, our mutational analyses indicated that such intranuclear condensation is mainly driven by electrostatic interactions in the form of a blocky pattern of opposite charges within the sequence of BRG1_C_, characteristic of polyampholytes. Indeed, IDRs containing patterned charge blocks have previously been shown to be an important driver for the condensation of other proteins through modulating their localization/partitioning, selective interactions and phase behavior (*22*, *36*, *37*), but not in the context of chromatin remodelers. As such, our findings not only broaden our general understanding of the condensation behaviors associated with various chromatin-based intranuclear processes (*38*–*40*), but also furnish a novel dimension to the “IDR grammar” that specifically governs the condensation and activities of SWI/SNF remodeling complexes (*e.g.* in addition to the role of uniformly distributed tyrosine residues known to drive the condensation of the ARID1A/B subunits (*14*).) More importantly, the sequence feature we unveiled is also in line with findings from recent analyses of the IDR grammar of nucleolar proteins, which identified D/E tracks and K-blocks + E-rich regions (ERRs) as distinct signatures for FC/DFC-localized proteins (*41*, *42*). Indeed, these IDR signatures are also extensively harbored within BRG1_C_ sequence (**Fig. S6C**), and are further corroborated by our finding that BRG1_C_ condensates selectively partition into the FC phase of the nucleolus.

The nucleolus is well-known to be a multiphase condensate organized in a hierarchical architecture, with distinct yet co-existing liquid phases each tailored for sequential steps in the ribosome biogenesis process; in particular, the boundary between the FC and DFC phases is the initiation site for this process, where rDNA repeats are located and transcribed into precursor rRNAs to be fluxed outward for further processing (*23*, *43*). Such a multiphase condensate is formed as a consequence of complex coacervation driven by a network of associative multivalent interactions, complemented by electrostatic interactions between various nucleolar components (*30*, *41*). As such, the patterned charge blocks we have unveiled in BRG1 could further contribute to the thermodynamic stability of such remodeler–nucleolus association. Moreover, while BRG1_C_ condensates are highly dynamic (as shown by FRAP), they remain stably nested within the FC phase unless perturbed by CX-5461- or ActD-induced nucleolar stress, where they relocate to the nucleolar periphery and form “nucleolar cap”-like structures, hence pointing to the key role played by Pol I-mediated rRNA transcription in modulating the architectural organization of BRG1_C_ condensates. Without such drug treatments, however, BRG1_C_ condensates within the same nucleolus do not fuse with each other across the DFC phase that surrounds them (**Fig. 3A**), in line with the previous observation that the DFC phase remains largely monodispersed even in the event of nucleolar coalescence (*44*). This suggests that such remodeler–nucleolus association driven by condensation is also structurally stable, while allowing for dynamic molecular exchange with the surrounding nucleoplasm.

Furthermore, our single-molecule tracking measurements found that BRG1_C_ within the nucleolar condensates exhibits substantially more constrained mobility (as manifested by diffusion coefficients and anisotropy measurements), indicative of a more viscous and crowded molecular environment. This, together with our observation that inhibiting rRNA transcription enhances the mobility of BRG1_C_ condensates, are in line with the earlier finding of the nucleolus as a complex viscoelastic fluid, with the FC phase being more gel-like due to the entanglement of nascent rRNAs, whereas the GC phase is more liquid-like following the maturation and folding of rRNAs into more compact ribosomal subunits (*30*). Moreover, the spatial densities of both the chromatin-binding events and the binding hotspots of BRG1_C_ are substantially increased within the nucleolar condensates for both transient and stable binding, in line with the FC phase being a more confined environment. In the temporal domain, the lifetimes of the binding hotspots are also significantly prolonged within the nucleolar condensates (especially so for the stable mode), as a consequence of adjoining more disparate stable binding events into a temporally longer hotspot; these hotspots likely correspond to sites posited for more sustained remodeling activity on the underlying rDNA, which is itself known to be a biomolecular condensate formed via polymer-polymer phase separation (*45*). Interestingly, both the frequency of binding events per unit time and the temporal modes of gap time between adjacent stable binding remain largely unchanged between inside and outside the condensates, indicating that spatial crowding does not impact foundational parameters of the temporal structure of BRG1’s binding dynamics. Overall, these findings demonstrate that in addition to the well-known effect of elevating the local concentrations of biomolecules, condensates can also enrich biomolecular interactions (such as chromatin-binding) in both space and time, by modulating both the spatial density as well as the temporal duration of the interactions involved at single-molecule level.

In light of such intricate interplay between BRG1 condensation, chromatin-binding dynamics and nucleolar architecture, we propose a mechanistic model for condensation-mediated remodeler–nucleolus coupling that integrates our findings from live-cell imaging, sequence pattern analysis, single-molecule tracking and correlative STAR/condensates mapping (**Fig. 6**). Leveraging the patterned charge blocks in its C-terminal IDRs, BRG1 uses condensation as an enrichment strategy to concentrate itself to the FC phase of the nucleolus, and undergoes an elevated level of chromatin-binding (particularly for stable binding). The higher spatial density of binding events as well as the prolonged lifetime of binding hotspots within the condensates potentiate BRG1 to more efficiently engage the underlying rDNA for sustained remodeling activity, and hence facilitate rRNA transcription within the nucleolus. The rRNAs transcribed in turn serve as a “nucleolar scaffold” to constrain the remodeler’s mobility, thereby helping to anchor and sequester it within the condensates. Such a spatiotemporal coupling could not only serve as a strategy for organizing remodeling activity in a locus-specific manner (in both physical and genomic spaces), but could also potentially synergize with the regulation of Pol I-mediated rRNA transcription (which is also known to occur via condensation (*31*)) to provide a coherent mechanistic paradigm for regulating ribosomal gene expression.

**Figure 6.**
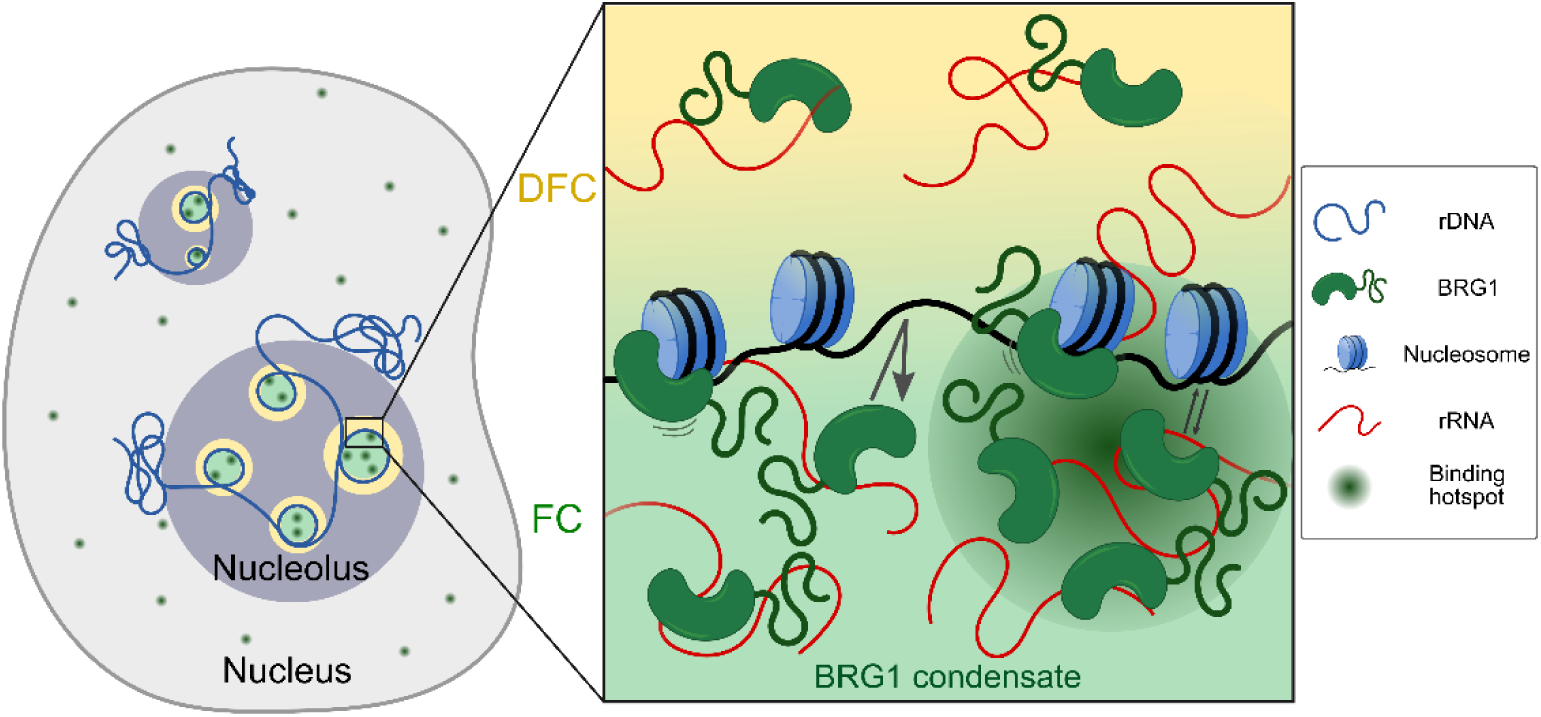
A model for condensation-orchestrated remodeler–nucleolus coupling to facilitate BRG1-mediated remodeling of rDNA. Driven by the patterned charge blocks in its IDR-rich C-terminus, BRG1 uses condensation as a strategy to enrich itself to the FC phase of the nucleolus and undergoes elevated chromatin-binding (particularly for stable binding). The higher spatial density of binding events as well as the prolonged lifetime of binding hotspots within the condensates potentiate it to more efficiently engage the rDNA (located at the FC/DFC boundary) for remodeling activity, hence facilitating rRNA transcription within the nucleolus. The rRNAs transcribed in turn serve as a “scaffold” to constrain BRG1’s mobility and sequester it within the nucleolar condensates.

Overall, the findings of this study shed critical insights into the functional importance of a distinctly patterned IDR in orchestrating the spatio-temporal organization and dynamics of a key chromatin remodeler in human. The condensation-mediated nucleolar coupling we have uncovered for BRG1 could potentially represent a generic mechanism used by remodelers to selectively regulate genome access, deepening our existing knowledge of the multi-faceted actions of chromatin remodelers in general (*1*, *46*). The live-cell correlative STAR/condensates mapping strategy devised in this study can also be applied to probe various other condensation-implicated intranuclear processes (*e.g.* transcription, splicing, DNA damage repair) at the single-molecule level, while paving the way for further interrogations into the mechanistic details of such condensation-mediated coupling (*e.g.* how the relative partitioning of the intranuclear BRG1 pool could be fine-tuned between the nucleolar condensates and the nucleoplasm for precise spatial control of remodeling activity). Moreover, our strategy could be correlatively integrated with single-molecule/super-resolution imaging of nucleosomes (*47*), euchromatin/heterochromatin (*48*), DNA loops (*49*), replication foci (*50*, *51*) and transcription factor binding (*52*, *53*) in the same cell to dissect the molecular interplay of BRG1 condensates with these key intranuclear structures. Lastly, combining our approach with proximal proteomics-based profiling (*54*, *55*) and rRNA sequencing (*56*, *57*) will help reveal the condensation-mediated interactome of BRG1 as well as its transcriptomic consequences, thereby elucidating the full functional implications of the unique remodeler–nucleolus coupling unveiled in this study.

## MATERIALS AND METHODS

### Constructs and cell lines generation

To generate constructs for mammalian expression of BRG1-mEmerald, the mEmerald gene sequence was amplified from pCS2+-mEmerald vector and cloned into pCS2+-hBRG1 vector (both gifts of Nicolas Plachta, University of Pennsylvania) via Gibson assembly using NEBuilder HIFI DNA Assembly Master Mix (E2621S, New England Biolabs).

To generate fusion constructs between BRG1_N_ or BRG1_C_ with mEmerald or HaloTag, pCS2+ vector was first linearized with restriction enzymes SpeI and BamHI (R3133 and R3136, New England Biolabs), and then assembled with the amplicons of mEmerald/HaloTag and BRG1_N_/BRG1_C_ via Gibson assembly. The mEmerald-BRG1_ΔC_ construct was subcloned by amplifying from the pCS2+-mEmerald-BRG1 construct, while the constructs of other mEmerald-BRG1_C_ truncation variants were subcloned by amplifying the desired region from the pCS2+-mEmerald-BRG1_C_ construct respectively.

To generate constructs of the various BRG1_C_ mutants, the respective gene sequences were designed and codon-optimized for DNA fragment synthesis by IDT. The synthesized DNA fragment with desired mutations was then ligated into the pCS2+ vector (with mEmerald at the N-terminus of the BRG1_C_ mutant) via Gibson assembly.

To generate construct for BRG1_C_ protein expression in bacterial cells, the sequence for 10xHis flanked with glycine linker was designed and purchased from IDT for annealing as dsDNA fragments, while the mEmerald and BRG1_C_ gene sequences were amplified from the pCS2+-mEmerald-BRG1_C_ construct. The three fragments were then ligated into pET28b protein expression vector via Gibson assembly, and amplified through DH5α cells for verification.

To generate eGFP fusion constructs for various nucleolar proteins, pEGFP gene sequence was first amplified by PCR from the pEGFP-C1-Fibrillarin vector (26673, Addgene). The respective gene sequences for NPM1, RPA194 and RPA43 were amplified from pPHR-mCherry-hNPM (181887, Addgene), p23-puro-EGFP-RPA194 (179237, Addgene) and pCA14-pHR-SFFV-GFP-RPA43 (P199445, Addgene) vectors respectively, and cloned into the pEGFP vector via Gibson assembly. On the other hand, vectors pEGFP-C1-Fibrillarin (26673, Addgene) and GFP-UBF (17656, Addgene) were used as purchased without additional modifications.

To generate CRISPR/Cas9 knock-in cell line that stably expresses mEmerald- or Halo-BRG1_C_ at endogenous level, a sgRNA (5’-TCTGGAGTGGACATCTTCAC-3’) targeting the transcription start site of BRG1 gene was designed using the CHOPCHOP platform (*58*), and cloned into the pCAG-spCas9-mCherry vector (a gift of Tan Meng How, Nanyang Technological University) by Gibson assembly, upon digestion with BbsI-HF restriction enzyme (R3539S, New England Biolabs). Homology arms flanking the knock-in site were amplified from the HeLa cell genome by PCR, and cloned into an empty PCR-linearized pCS2+ vector together with the knock-in sequence for mEmerald- or Halo-BRG1_C_-P2A via Gibson assembly. Then, 1 µg of the CRIPSR/Cas9 plasmid and 1 µg of the homology-directed repair template (as circular plasmid) per well were co-transfected into Hela cells seeded in a 6-well plate. Positive cells were enriched by FACS detected at 488 nm (for mEmerald) or 561 nm (for HaloTag labelled with TMR); single clones were then isolated by sorting positive populations into a 96-well plate, and subjected to further screening by PCR and genomic validation.

All constructs generated were validated by sequencing before transfection or transformation. All primers used to generate the constructs are listed in **Table S1**.

### Cell culture and transfection

HeLa cell line (ATCC CCL-2, a gift from Thorsten Wohland, National University of Singapore) was maintained in HyClone Dulbecco’s modified Eagle’s medium (DMEM) with high glucose (SH30022.LS, Cytiva) supplemented with 10% (v/v) fetal bovine serum (FBS, A5256701, Gibco) and 1% (v/v) penicillin-streptomycin (Pen-Strep, 15140122, Gibco), and cultured at 37 °C in a humidified incubator with 5% CO_2_. For live-cell and immunofluorescence imaging, cells were first seeded on a glass-bottom dish with a 10-mm microwell and No. 1.5 coverglass (P35G-1.5-10-C, MatTek) at ∼80% confluency one day prior to transfection or fixation. Cells were then transfected overnight with 400-600 ng of plasmid DNA using Lipofectamine 3000 (L3000-015, Thermo Fisher Scientific) according to manufacturer’s protocol.

### Immunofluorescence

Hela cells were fixed 24 hours post-seeding or post-transfection with 3% (w/v) paraformaldehyde and 0.1% (w/v) glutaraldehyde in 1x phosphate buffered saline (PBS) for 10 minutes at room temperature (RT). After two washes with 1x PBS, the cells were first blocked for 1 hour with blocking buffer (5% (w/v) bovine serum albumin (BSA, A3059, Sigma-Aldrich) and 0.5% (v/v) Triton X-100 (A16046, Thermo Fisher Scientific) in PBS) at RT, and then incubated with the respective primary antibody (1:500 mouse anti-NPM1 (32-5200, Invitrogen) or 1:1000 rabbit anti-fibrillarin (ab5821, Abcam)) diluted in blocking buffer for 1 hour at RT with gentle agitation. The sample was then washed three times with washing buffer (0.5% (w/v) BSA and 0.05% (v/v) Triton X-100 in PBS) at RT, followed by incubation with the respective Alexa Fluor® 647-labeled secondary antibody (1:1000 goat anti-mouse IgG (ab150115, Abcam) or 1:1000 donkey anti-rabbit IgG (ab150075, Abcam)) diluted in blocking buffer for 40 minutes at RT with gentle agitation. Upon washing three times with 1x PBS, the sample was then incubated with 1 µg/mL DAPI (MBD0015, Sigma) in PBS for 10 minutes at RT, and washed three times with 1x PBS before imaging.

### Confocal microscopy

Live or fixed HeLa cells (seeded at the appropriate confluency on either glass-bottom dish (P35G-1.5-10-C, MatTek) or Nunc™ Lab-Tek™ II chambered coverglass (155382, Thermo Fisher Scientific) were imaged on a Nikon Eclipse Ti inverted microscope, equipped with 4 diode laser lines (at excitation wavelengths 405 nm (60 mW, LuxX® 405-60, Omicron), 488 nm (200 mW, LuxX® 488-200, Omicron), 561 nm (150 mW, OBIS™ 1280720, Coherent) and 647 nm (200 mW, LuxX® 638-200, Omicron)), a Nikon C2si confocal scanhead, and a Okolab microscope stage-top chamber maintained in a humidified environment at 37 °C with 5% CO_2_ supply. Prior to imaging, cells were incubated with Hoechst 33342 (NucBlue™, R37605, Life Technologies) in imaging medium consisting of phenol red-free DMEM (21063029, Life Technologies) for at least 10 minutes at 37 °C to stain the DNA/cell nucleus. Confocal images were acquired with a 60x, NA 1.20 water immersion objective (CFI Plan Apo VC 60XC WI, Nikon) at a scanning rate of ∼4.8 µs/pixel and 1–4% laser power, using the Nikon NIS-Element AR software.

### Protein expression and purification

To express mEmerald-10xHis-BRG1_C_ protein in *E. coli* system, the expression construct was transformed (according to manufacturer’s instruction) into Rosetta^TM^ (DE3) competent cells (70-954-4, Merck Millipore) cultivated in LB medium (for seed culture) or BRM medium (for scale-up culture). The bacterial culture was grown at 37 °C until a OD_600_ of ∼ 0.6 was reached, and then chilled on ice for 10-15 minutes before being induced with 0.5 mM isopropyl β-d-1-thiogalactopyranoside (IPTG) (GS-25, Alkali Scientific). The induced culture was grown at 30 °C for 6–8 hours, before harvesting the cells with centrifugation at 4,000 rpm for 15 minutes at 4 °C. To purify the mEmerald-10xHis-BRG1_C_ protein, the cell pellet obtained was resuspended and lysed in lysis buffer (300 mM NaCl, 10% glycerol, and 2 mM CHAPS in 50 mM Tris at pH 8.0) with sonication, and centrifuged at 14,000 rpm for 20 minutes at 4 °C. The supernatant collected was first subjected to Ni-NTA purification (30210, Qiagen), eluted with 250 mM imidazole in elution buffer (lysis buffer supplemented with 2 mM β-mercaptoethanol), and further purified with a HiTrap Q HP anion exchange chromatography column (17115401, Cytiva) on a ÄKTA go^TM^ FPLC system (Cytiva). The protein was eluted with a linear gradient from 20% to 50% of buffer B containing 1 M NaCl, and the eluent fractions were fractionated and analyzed with SDS-PAGE. The selected fractions were pooled and concentrated to ∼1 mL for subsequent purification with size exclusion chromatography on a Superdex^TM^ 200 10/300 GL column (28990944, Cytiva), and the eluent fractions were quality-checked with SDS-PAGE. The desired fractions were then concentrated again using a 50 kDa MWCO Amicon® Ultra centrifugal filter (UFC8050, Millipore) at 4 °C, and aliquoted into small volumes for flash-freezing and long-term storage at −80 °C.

### *In vitro* droplet assay

*In vitro* droplet assay was set up in a 10 µL reaction system on ice for each condition tested. The purified and concentrated mEmerald-10xHis-BRG1_C_ protein was diluted in 50 mM Tris buffer at pH 8.0 to the desired concentration. Appropriate amounts of NaCl (from a 4 M stock) and/or PEG-8000 (P2139, Sigma-Aldrich, from a 50% (w/v) stock) were then added to reach the desired final concentration to be tested. Upon incubation for at least 10 minutes, the mixture was then added onto a No. 1.5 coverglass and imaged under confocal microscopy after 5 minutes of system stabilization and equilibration to RT, focusing on droplets at the surface of the coverglass. The acquired images were then analyzed with Fiji (*59*).

### Fluorescence recovery after photobleaching (FRAP)

HeLa cells seeded at the appropriate confluency were transfected with pCS2+-mEmerald-BRG1_C_ construct one day prior to FRAP measurement. The cells were imaged using. FRAP measurement was carried out on a Olympus FLUOVIEW FV3000 confocal microscope, equipped with 20 mW 488-nm laser (OBIS™ 1178767, Coherent), a Okolab stage-top chamber maintained in a humidified environment at 37 °C with 5% CO_2_ supply, and a 60x water immersion objective (UPLSAPO60XW, Olympus). The FRAP bleaching and acquisition workflows were pre-programed using FV31S-SW Viewer software. Each region of interest (ROI) was first bleached at 7% laser power for 0.2 s, and then continuously imaged at 2% laser power to monitor the fluorescence recovery. The images acquired were then analyzed with cellSens software to obtain the raw bleaching and recovery intensity data from each ROI, the associated time stamps and the various fitted parameters.

### 1,6-Hexanediol/drug treatment

For 1,6-hexanediol (1,6-HD) treatment, HeLa cells was transfected with mEmerald-BRG1_C_ construct one day prior to treatment, and incubated with 1,6-HD (240117, Sigma-Aldrich) at the respective concentration (in w/v) diluted from a 20% (w/v) stock in DMEM complete medium. For CX-5461 or actinomycin D (ActD) treatment, HeLa cells stably expressing mEmerald-BRG1_C_ were seeded two days prior to treatment, and transfected with pCS2+-mCherry-NPM1 construct one day prior to treatment. Upon replacing with fresh DMEM complete medium, cells were treated with either 500 nM CX-5461 (HY-13323, MedChemExpress) for 4 hours or 0.1 µg/mL ActD (A9415, Sigma-Aldrich) for 1 hour at 37 °C, followed by incubation with Hoechst 33342 for at least 10 minutes at 37 °C to stain the DNA/cell nucleus. After washing two times with DMEM complete medium and replacing with fresh DMEM complete medium, cells were allowed to recover over time, while being imaged with confocal microscopy. For suberoylanilide hydroxamic acid (SAHA) treatment, HeLa cells stably expressing Halo-BRG1_C_ were seeded one day prior to treatment, and incubated with 5 µM SAHA (SML0061, Sigma-Aldrich) for 24 hours prior to SMT imaging.

### Bioinformatic analyses

Sequence-ensemble relationships for BRG1, BRG1_N_ and BRG1_C_ (including net charge per residue, fraction of charged residues, hydrophobicity, κ values, and compositional bias) were calculated using the CIDER platform (*21*) and classified in the form of diagram of states. Each sequence was classified as a foldable protein or an intrinsically disordered protein (IDP) using the Uversky plot based on its mean hydrophobicity and mean net charge (*60*), generated with CIDER. Non-random sequence patterning features of BRG1, BRG1_N_, BRG1_C_ and BRG1_C_^CS^ were calculated using NARDINI (*22*). To do so, the 20 canonical amino acids were grouped into eight distinct categories (polar, hydrophobic, positively charged, negatively charged, aromatic, alanine, proline and glycine), and a null model of 10^5^ randomly scrambled sequences with fixed amino acid composition from each input sequence was generated to calculate the z-scores; a z-score of >0 indicates the residue groups are de-mixed into blocks in the linear sequence, while a z-score of < 0 indicates that the residues are well-mixed throughout the linear sequence. Lastly, BRG1 structure prediction was performed using the AlphaFold DB platform (*61*) (https://alphafold.ebi.ac.uk).

### Live-cell single-molecule tracking (SMT)

HeLa cells stably expressing Halo-BRG1_C_ were seeded at the appropriate confluency one day prior to SMT measurement, and incubated on the day with 2.5 nM Janelia Fluor® 549 (JF549) Halo-tag® ligand (GA1110, Promega) for 15 minutes at 37 °C, followed by three times of washing with DMEM complete medium for 10 minutes each to remove the excess dyes. Prior to SMT acquisition, the cells were incubated with Hoechst 33342 in imaging medium consisting of phenol red-free DMEM for at least 10 minutes at 37 °C to stain the DNA/cell nucleus. Cells were imaged live on a Nikon Eclipse Ti2-E inverted microscope equipped with 405-nm (100 mW, OBIS™ 1284371, Coherent) and 561-nm (150 mW, OBIS™ 1280720) laser lines, a 100x, NA 1.49 oil objective (SR HP Apo, Nikon), and a Okolab microscope stage-top chamber maintained in a humidified environment at 37 °C with 5% CO_2_ supply. Highly inclined and laminated optical sheet (HILO) illumination (*62*) was used to suppress intranuclear background for improved signal-to-background ratio in SMT data. Before and after each SMT acquisition, an image of the cell nucleus was acquired under 405-nm excitation at low power for isolating only intranuclear trajectories in subsequent image analysis. A 20-frame image stack (at 50 ms per frame) of BRG1_C_ condensates was also acquired for each cell at 561 nm before each SMT acquisition for subsequent correlative analysis. After a brief illumination to bleach excessive background signal, SMT acquisition was commenced and images of single BRG1_C_ molecules were continuously acquired at 561 nm using the Nikon NIS-Element AR software, for 10,000 frames at 5.5 ms per frame and a power density of ∼0.4 kW/cm^2^ (for fast tracking) or for 2,000 frames at 300 ms per frame and a power density of ∼0.02 kW/cm^2^ (for slow tracking).

When quantifying the impact of drug treatments (e.g. CX-5461 and SAHA) on BRG1_C_ dynamics, every SMT dataset for each drug treatment was accompanied by a control dataset (without drug treatment) acquired on the same day on identically prepared cell samples. Around 15 cells per treatment condition were measured per day, and the distribution of each dynamic parameter measured was normalized against the median value of the control distribution.

### SMT data analysis

SMT data analysis was performed as previously described (*17*) with modifications. Briefly, each frame of a SMT movie was first denoised with a wavelet filter (*63*), and the 2D coordinates for individual localizations of BRG1_C_ molecules were then extracted with a radial gradient-based algorithm (*64*) using a custom-written plugin operating in Fiji. All subsequent analyses were performed in Matlab (MathWorks) using custom-written scripts. Firstly, trajectories were constructed from extracted localizations using the TrackMate plugin on Matlab (*65*). Specifically, each trajectory was required to consist of at least 3 or 2 consecutive displacements for fast- and slow-tracking respectively, with a maximum allowed displacement between adjacent frames set at 640 nm and 280 nm for fast- and slow-tracking, respectively; a maximum of 2 consecutive missed frames was allowed for both modes. From fast-tracking trajectories, the diffusion coefficient (*D*) and the corresponding molecular fraction (*f*) for each mobility mode of BRG1_C_ were extracted by fitting the normalized displacement histogram obtained from all trajectories with a three-state model (*29*):

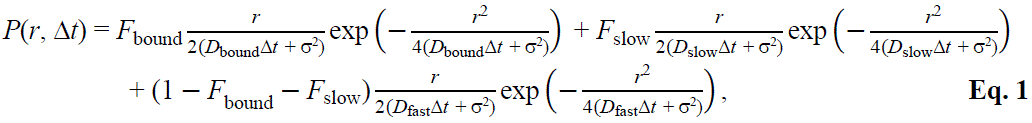

where the subscripts “fast”, “slow” and “bound” denote the fast-diffusion, slow diffusion and bound modes respectively, while σ denotes the localization uncertainty (15 nm in our case). From slow-tracking trajectories, binding events were first unambiguously identified by checking all possible sub-trajectories and picking those that are confined within a circular area with an optimized size. Only binding events with at least 4 consecutive displacements were chosen to avoid the inclusion of trajectories from slowly diffusing molecules that might kinetically resemble bound molecules. To calculate binding parameters, the survival probability for a molecule to remain bound after a certain time *t* was first calculated, and the resulting distribution was analyzed using GRID (*33*) to determine the number of binding modes involved. The survival probability distribution was then fitted with a multi-exponential function based on the number of binding modes identified from GRID to obtain the residence time (τ) and the corresponding molecular fraction (*f*) for each mode.

### Correlative STAR/condensates mapping

STep Accumulation Reconstruction (STAR) mapping analysis was performed as previously described (*17*) with modifications. Briefly, a grid with a pixel size determined by the number of displacements detected and the associated localization precision (40 nm in our case) was first defined. The centroid position of each displacement in a binding trajectory of BRG1_C_ was then mapped onto the grid, with the brightness of each pixel corresponding to the number of displacements that fall into that pixel. An intranuclear STAR map for binding can then be generated by plotting the density of displacements for each pixel, which can then be smoothened by applying a Gaussian filter (with σ = 40 nm); further classifying the binding events based on the duration of residence time then yields a binned map that can be displayed in RGB format. To detect the spatial clustering of binding events into binding hotspots, a distance-based analysis was performed using pairwise distances between binding events. Similarly, a STAR map for diffusion can be generated using sub-trajectories that correspond to diffusion of BRG1_C_. Finally, to correlate the binding or diffusion map with intranuclear condensates distribution, a wide-field image of the cell was acquired before each SMT acquisition to generate a binary mask for isolating the cell nucleus and condensates in diffusion or binding maps; the intranuclear diffusion or binding trajectories can then be classified into either inside or outside the condensates accordingly.

### Electrophoretic mobility shift assay

A complementary pair of 21-base DNA fragments from the Widom 601 sequence (CTCAATTGGTCGTAGACAGCT) was purchased from IDT and annealed as dsDNA. The dsDNA was added to a 10 µM solution of purified mEmerald-10xHis-BRG1_C_ protein (in 50 mM Tris (pH 8.0), 150 mM NaCl and 20% glycerol) to a final concentration of 0.5 µM; serial dilutions were then performed to achieve the desired [protein]:[DNA] ratios of 0:1, 1:1, 2:1, 4:1, 8:1, and 12:1, respectively. The samples were incubated on ice for 20 minutes before loading onto a 1% TBE-agarose gel, and electrophoresed at 70–80 V for 30 minutes at 4 °C. As control, a dsDNA-only sample at concentrations ranging from 31.25 nM to 1000 nM was also run side-by-side.

### Statistical analysis

All biological replicate experiments were derived from at least three independent measurements conducted on different days. For box-and-whisker plots, the median, 25^th^/75^th^ percentiles (box) and 5^th^/95^th^ percentiles (whiskers) are shown. Statistical significance was assessed using unpaired Student’s *t*-test: *: *p* < 0.05; **: *p* < 0.01; ***: *p* < 0.001; NS: not significant.

## ACKNOWLEDGEMENTS

We thank Nicolas Plachta for the gift of the pCS2+-hBRG1 and pCS2+-mEmerald plasmids, Tan Meng How for the gift of the pCAG-spCas9-mCherry plasmid, Thorsten Wohland for the gift of HeLa CCL-2 cell line, Siyi Chen for helpful discussion on cloning, Goran Biukovic for early effort in establishing the protein purification workflow, Min Luo for access to the ÄKTA go^TM^ FPLC system, Joel Ng and Shifeng Xue for advice on protein expression/purification, and Thorsten Wohland and Pakorn Kanchanawong for critiques and helpful suggestions.

## FUNDING

This work was supported by the Ministry of Education of Singapore Academic Research Fund Tier 2 Grant (T2EP30222-0038) and Tier 3 Grant (MOET32020-0001), as well as the National University of Singapore Presidential Young Professorship Start-up Fund to Z.W.Z.

## AUTHOR CONTRIBUTIONS

Conceptualization: ZWZ, WSN; Methodology: WSN, ZWZ; Software: WE; Investigation: WSN; Formal analysis: WSN, WE; Data curation: WE, WSN; Writing-Original Draft: WSN, ZWZ; Writing-Review & Editing: All authors; Visualization: WSN, WE; Supervision: ZWZ; Funding acquisition: ZWZ.

## COMPETING INTERESTS

The authors declare no competing interests.

## DATA AND MATERIALS AVAILABILITY

All data needed to evaluate the conclusions are presented in the paper and/or the Supplementary Materials. Due to their large size, the raw SMT data are available from the corresponding author upon reasonable request. All codes used in this study are available from GitHub at https://github.com/wilengl/Nat_Comm_Codes/.

## Supplementary Materials

**This PDF file includes:**

Figures S1 to S13

Table S1

Supplementary References 1 to 5

**Figure S1.**
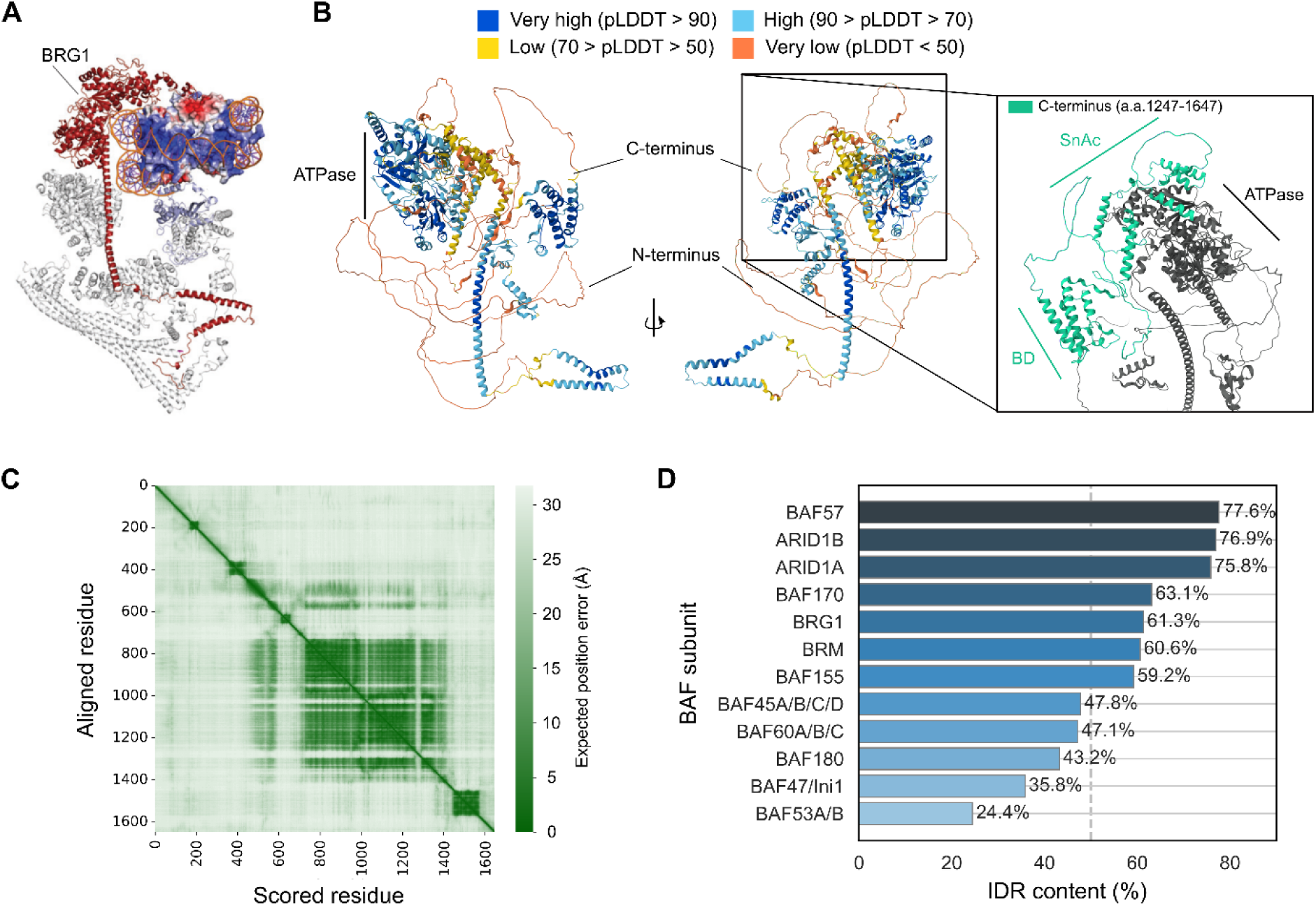
BRG1 and many other key SWI/SNF subunits contain high levels of IDRs in their structures. **(A)** A partially resolved cryo-EM structure of human BAF complex (in cartoon representation), with the resolved parts of the BRG1 subunit highlighted in red; the nucleosome is shown in electrostatic surface representation. Adapted from (*1*). **(B)** Two different views of the full structure of BRG1 predicted by AlphaFold DB (with each residue color-coded according to its associated predicted local distance difference test (pLDDT) score), showing the presence of extensive IDRs. Inset shows a zoomed-in view of the C-terminus (*i.e.* residues 1247−1647) in mint green, while the rest of the protein is shown in grey. **(C)** Predicted aligned error (PAE) heatmap of each residue in AlphaFold DB-predicted structure of BRG1, with the shade of color at each coordinate (*x*, *y*) indicating the expected distance error (in Å) at residue *x*’s position when the predicted and the true structures are aligned on residue *y*. **(D)** High levels of IDRs are present in many of the key SWI/SNF remodeler subunits. Data adapted from (*2*).

**Figure S2.**
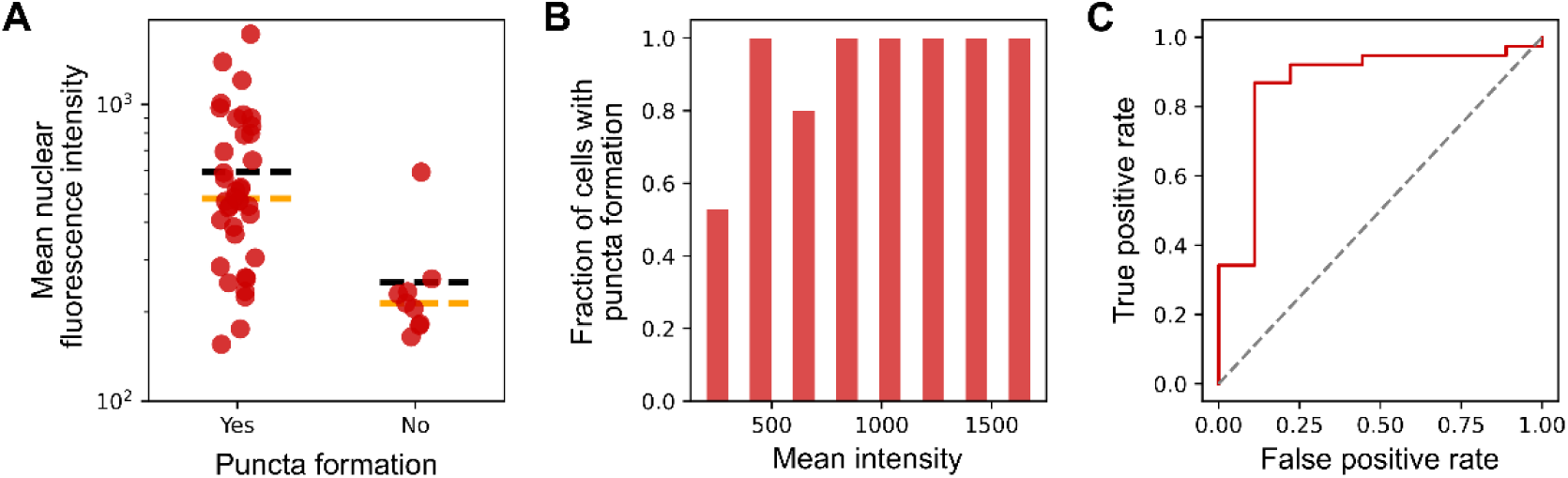
Quantitative characterization of intranuclear BRG1 condensates formation. **(A)** Scatter plot of mean nuclear fluorescence intensity of live Hela cells ectopically expressing mEmerald-BRG1, sorted according to whether they exhibit BRG1 puncta formation or not. **(B)** Histogram of the fraction of cells exhibiting BRG1 puncta formation as a function of mean nuclear fluorescence intensity. **(C)** Receiver Operating Characteristic (ROC) analysis indicates that mean nuclear fluorescence intensity can serve as a reliable predictor for BRG1 puncta formation. In **A**, each dot denotes a single cell. *n* = 47 cells.

**Figure S3.**
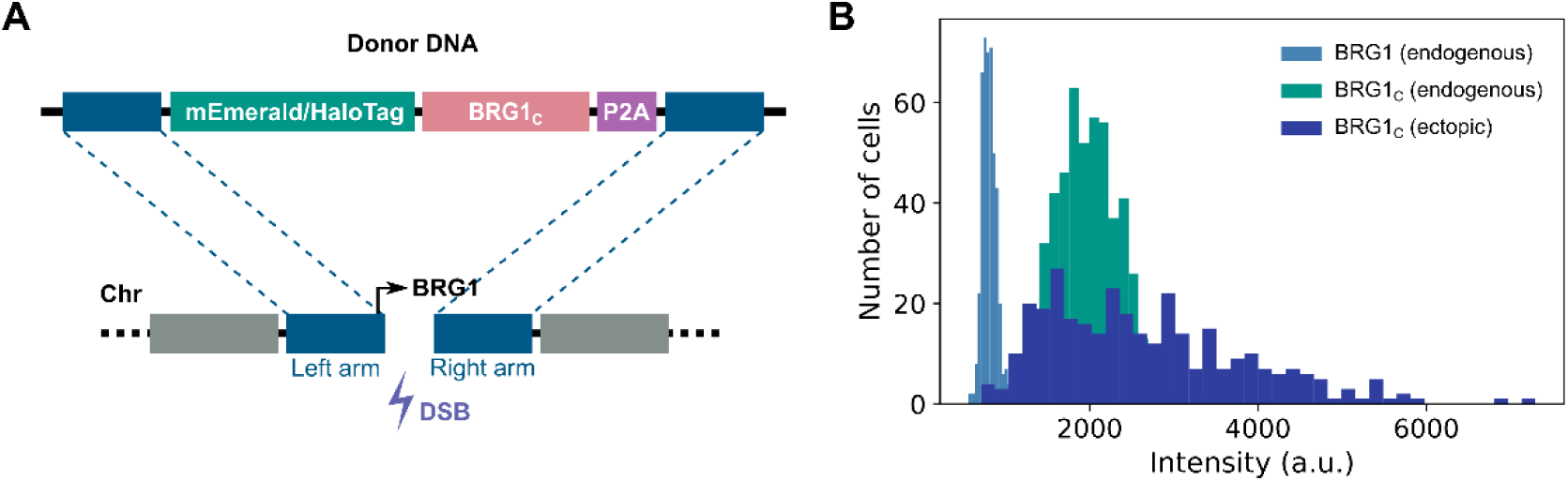
Quantitative comparison of ectopic and endogenous expression levels of BRG1_C_. **(A)** Schematic for the generation of CRISPR-Cas9 knock-in Hela cell lines that endogenously express mEmerald- or Halo-tagged BRG1_C_. **(B)** Histogram of the mean nuclear fluorescence intensity of each cell from the CRISPR-Cas9 knock-in cell lines expressing either BRG1-Halo (*3*) or Halo-BRG1_C_ at endogenous levels, in comparison with cells ectopically expressing Halo-BRG1_C_. Only cells with expression levels that fall within the overlapping region between the endogenous and ectopic expression level distributions for Halo-BRG1_C_ were selected for subsequent characterizations to ensure consistency and comparability. For **B**, *n* = 476, 505 and 328 cells for BRG1-Halo, Halo-BRG1_C_ (endogenous) and Halo-BRG1_C_ (ectopic), respectively.

**Figure S4.**
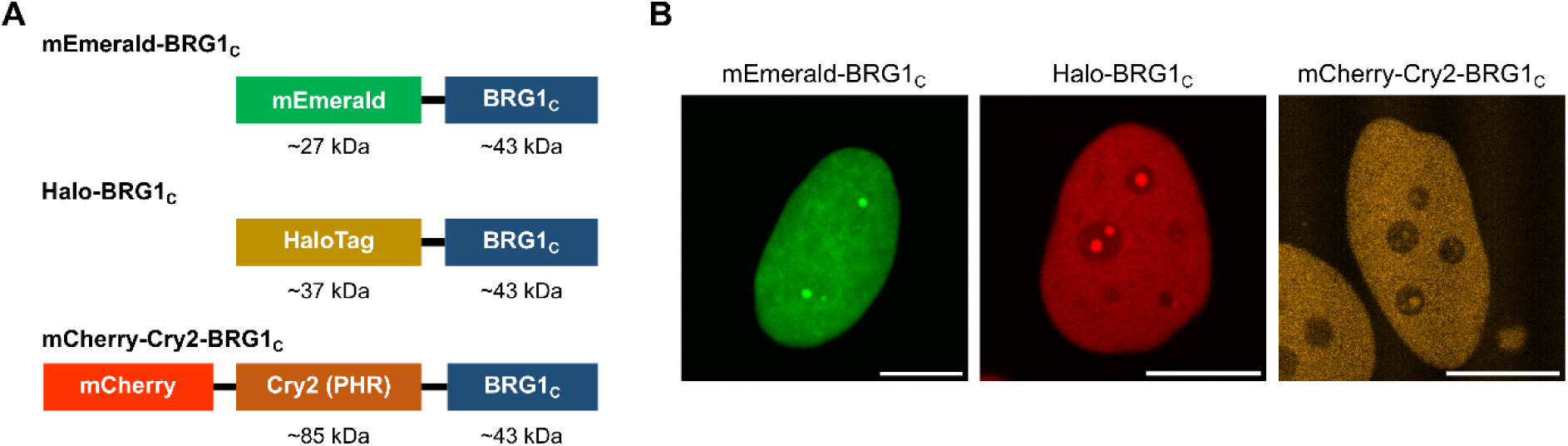
BRG1_C_ condensate formation is independent of the fluorescent protein or tag to which it is fused. **(A)** Schematics of different BRG1_C_ fusions constructed with fluorescent protein or self-labeling tag of different molecular weights. **(B)** Confocal images of live HeLa cells expressing either mEmerald-BRG1_C_, Halo-BRG1_C_ (labeled with 5 nM Janelia Fluor 646 Halo-tag ligand) or mCherry-Cry2-BRG1_C_, exhibiting similar puncta formation. Scale bar for **B**: 10 µm.

**Figure S5.**
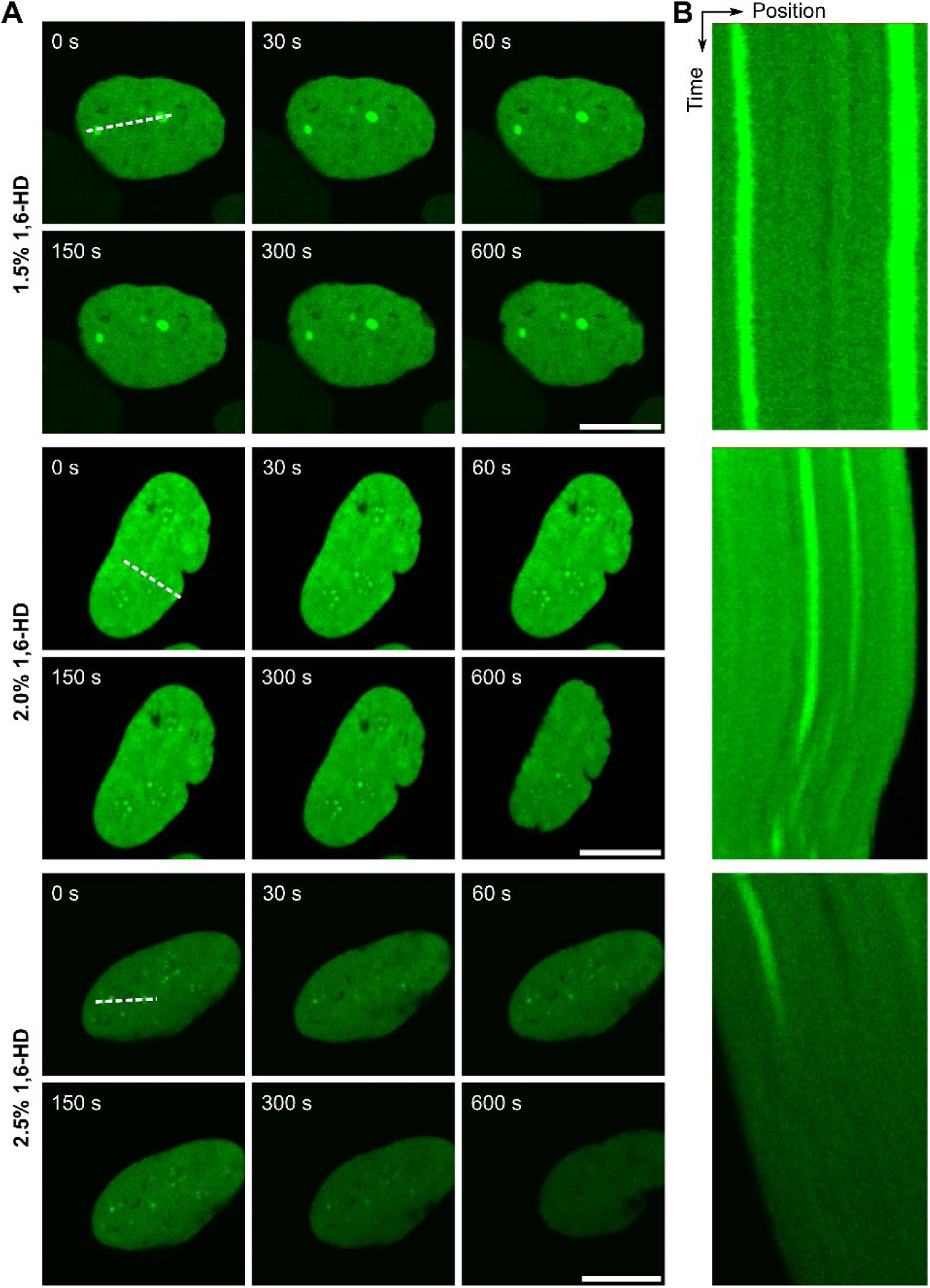
Intranuclear BRG1_C_ condensates are largely insensitive to 1,6-HD treatment across a range of concentrations. **(A)** Confocal images of mEmerald-BRG1_C_ condensates in live HeLa cells at different time points across a period of 600 s upon treatment with either 1.5% (top), 2.0% (middle) or 2.5% (bottom) 1,6-HD (all in w/v). **(B)** Kymographs of fluorescence intensities across the white dotted lines in **(A)** across a period of 600 s upon 1,6-HD treatment. The disappearance of the BRG1_C_ condensate signals in the latter parts of the kymographs for 2.0% and 2.5% 1,6-HD treatment was due to the gradual shrinking and drifting away of the cell from the focal plane, rather than the dissolution of the condensates. Scale bar for **A**: 10 µm.

**Figure S6.**
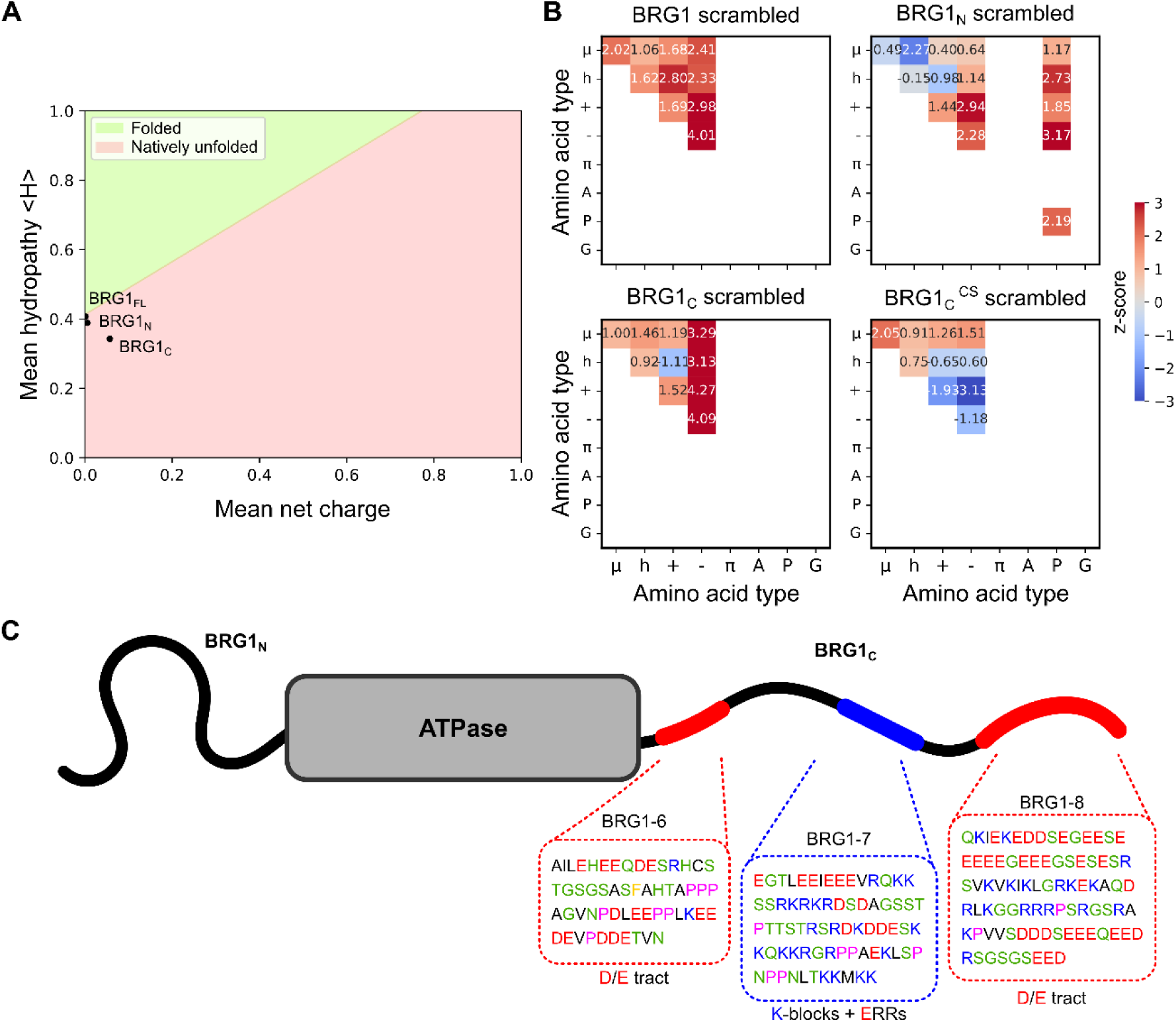
Bioinformatic analyses indicate that BRG1_C_ is a strong polyampholyte enriched in patterned charge blocks. **(A)** Uversky plot of the disorder state generated with CIDER based on protein net charge and hydropathy allocates BRG1_C_ to the natively unfolded region, while BRG1_N_ and BRG1_FL_ fall on the border between natively unfolded and folded proteins. **(B)** Z-score matrices of sequence patterning parameters calculated by NARDINI for null-scrambled models of BRG1, BRG1_N_ and BRG1_C_ (both wild-type and charge-scrambled mutant). Residues are grouped according to polar or μ (S, T, N, Q, C, H), hydrophobic or h (I, L, M, V), positively charged or + (R, K), negatively charged or − (E, D), aromatic or π (F, W, Y), alanine (A), proline (P), and glycine (G). **(C)** Schematic of BRG1 sequences showing the presence of charged D/E tracts and K-block + E-rich regions (ERRs) within its C-terminus, as shown by King *et al.* (*4*); residues are colored according to positively (blue) or negatively (red) charged, polar (green), hydrophobic (black) and proline (pink), respectively.

**Figure S7.**
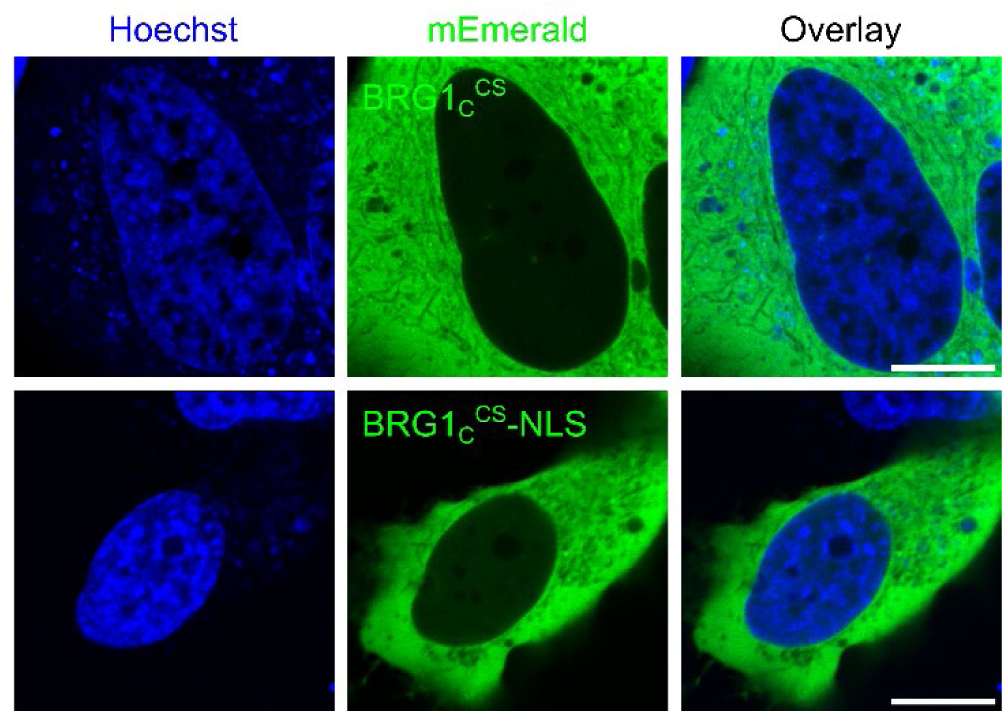
Intranuclear localization and condensate formation of BRG1_C_ are abolished in charge-scrambled mutants. Confocal images of live HeLa cells expressing the mEmerald-BRG1_C_^CS^ mutant either without (top row) or with (bottom row) a fused nuclear localization sequence (NLS); cell nucleus was co-stained with Hoechst. Both the intranuclear localization and condensate formation of BRG1_C_ are abolished in these charge-scrambled mutants. Scale bar: 10 µm.

**Figure S8.**
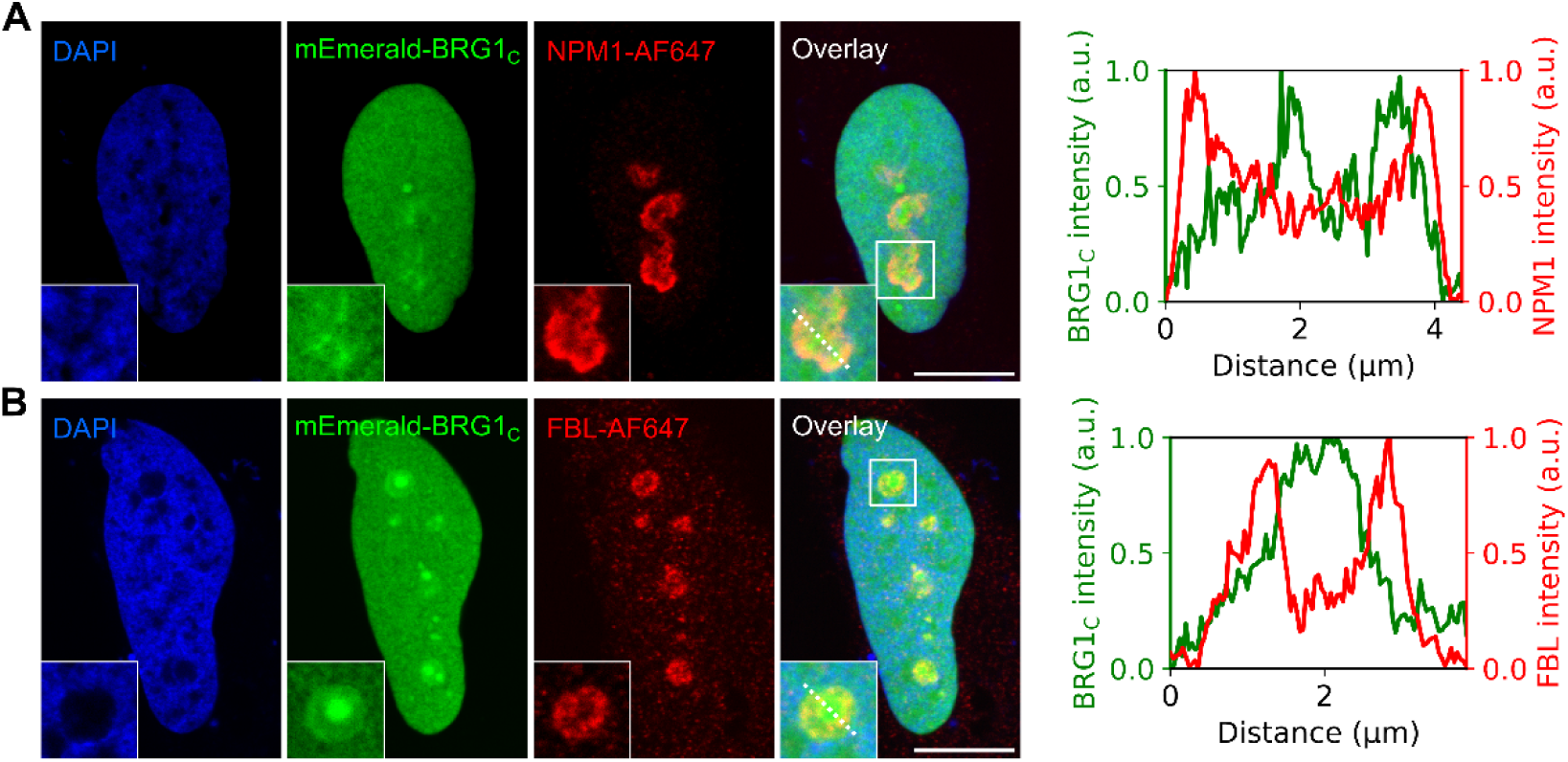
Colocalization of BRG1_C_ condensates with the FC phase of nucleolus validated with immunofluorescence imaging. (**A**–**B**) Confocal images of Hela cells ectopically expressing mEmerald-BRG1_C_ and immunofluorescently labeled with key nucleolar markers: NPM1 of granular component (GC) (**A**) and fibrillarin (FBL) of dense fibrillar component (DFC) (**B**). Cell nucleus was co-stained with DAPI. Insets show zoomed-in regions in the overlay images delineated by white boxes, with the fluorescence intensity changes across the white dotted lines plotted on the right. The lack of colocalization between BRG1_C_ condensates and either the GC or the DFC phase corroborates the observation reported in Fig. 3. Scale bar: 10 µm.

**Figure S9.**
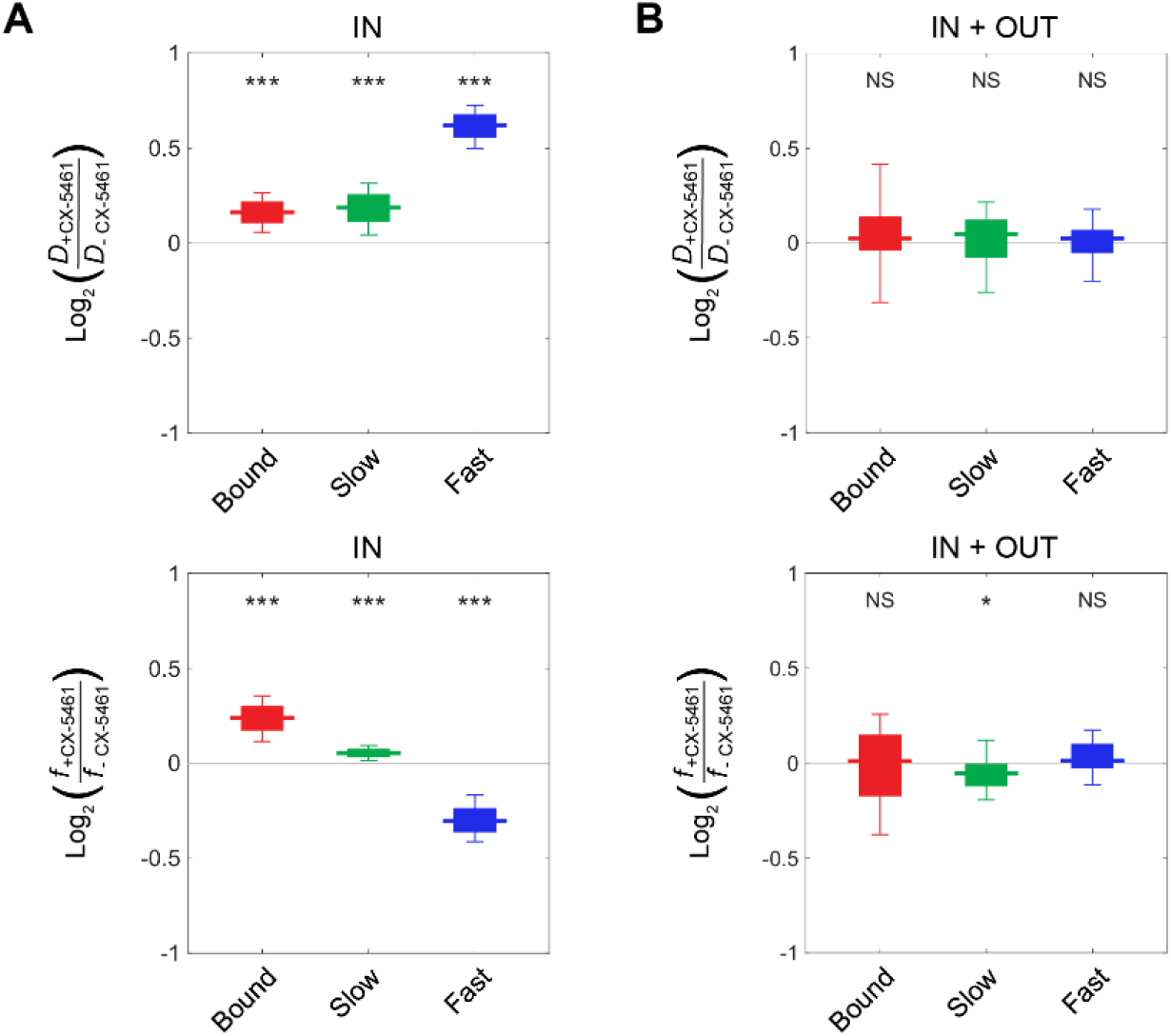
The effect of CX-5461 treatment on BRG1_C_ mobility is no longer discernible upon pooling all SMT trajectories from both inside and outside nucleolar condensates. (**A**–**B**) Box-and-whisker plots of fold changes (on a log2 scale), upon CX-5461 treatment, in *D* (top) and *f* (bottom) associated with each of the three modes of BRG1_C_ mobility, using only trajectories from inside the BRG1_C_ condensates **(A)** (*i.e.* Fig. 4H) or by combining all trajectories from both inside and outside the condensates **(B)**. *n* = 28 and 30 cells for with and without CX-5461 treatment, respectively.

**Figure S10.**
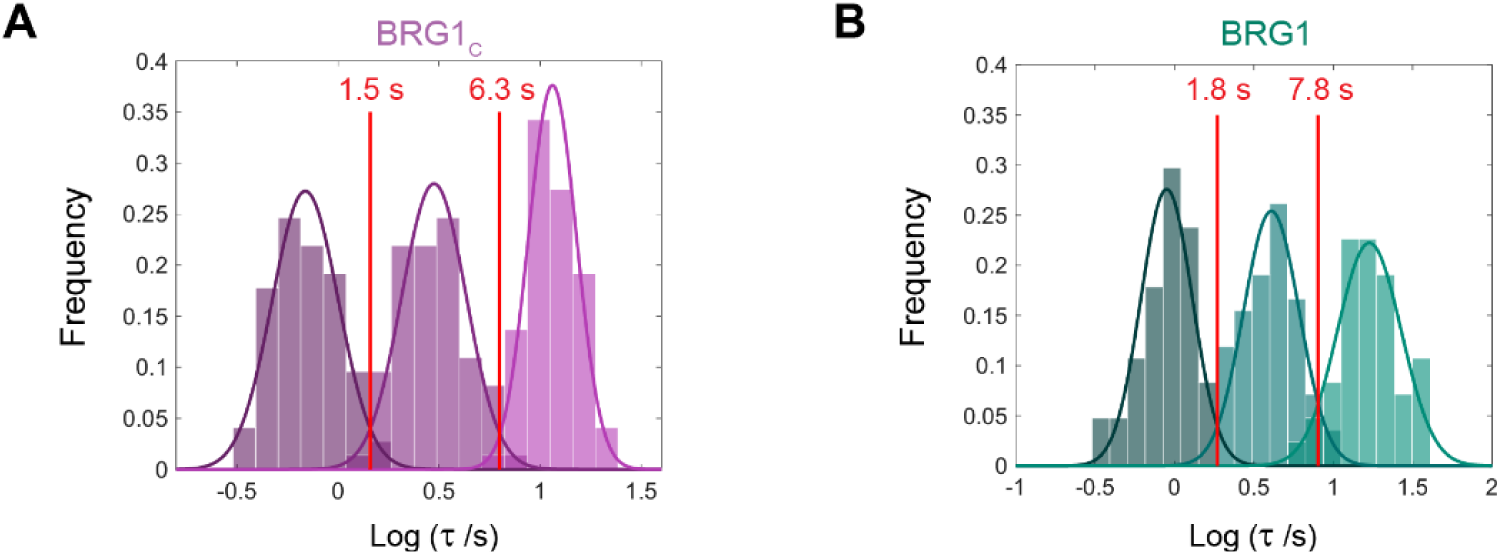
BRG1_C_ exhibits three temporally distinct modes of chromatin-binding with similar residence times to full-length BRG1. (**A**–**B**) Residence times distribution for the three chromatin-binding modes of BRG1_C_ **(A)** and full-length BRG1 **(B)**, each fitted with a triple-Gaussian fit; the temporal ranges associated with each of the three modes are demarcated by red lines. *n* = 73 cells for **A** and 84 cells for **B**. Data in **B** adapted from (*3*).

**Figure S11.**
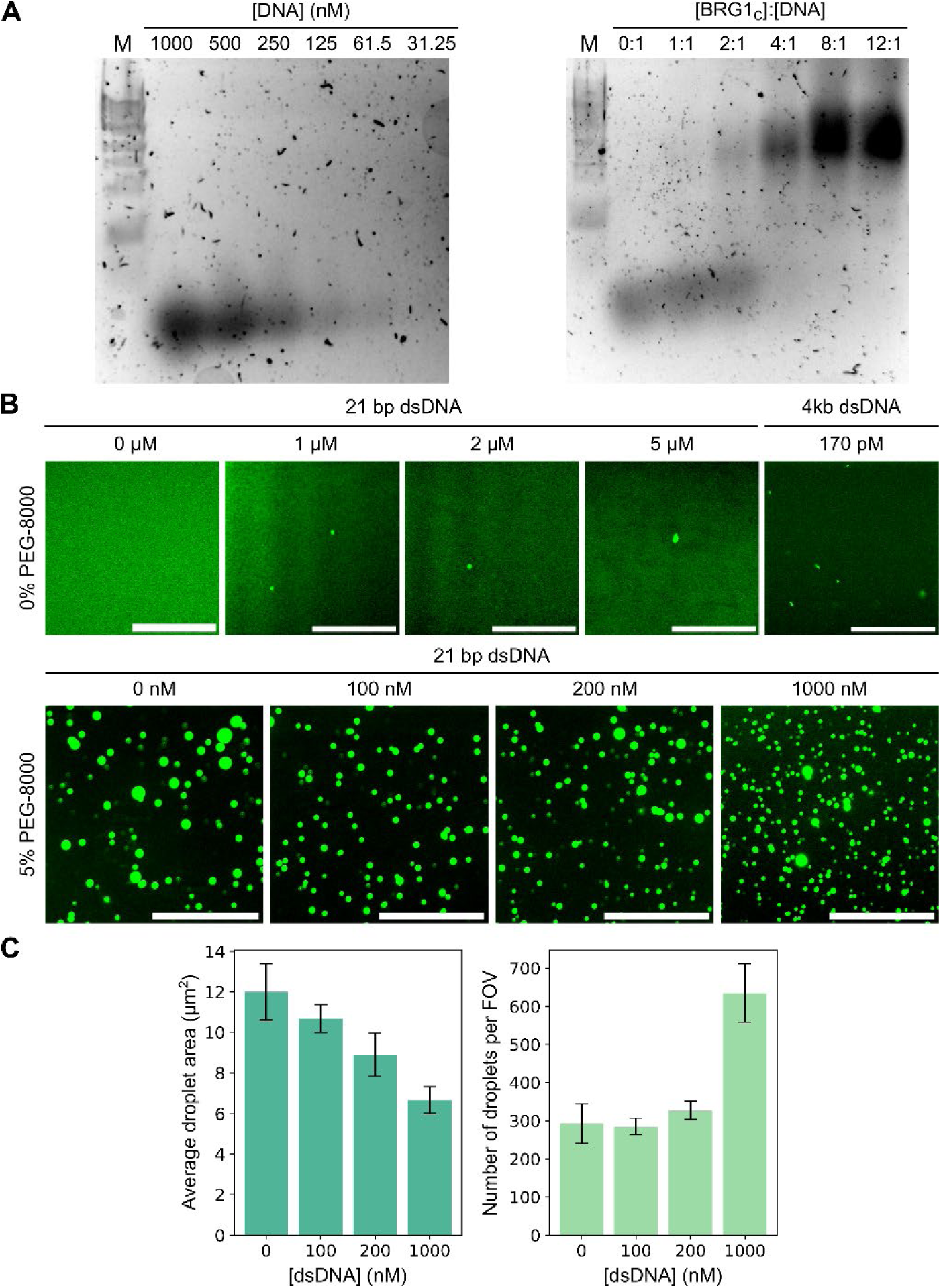
BRG1_C_ condensation *in vitro* is enhanced by nucleosomal DNA. **(A)** Electrophoretic mobility shifty assay (EMSA) on purified mEmerald-10xHis-BRG1_C_ in the presence of 0.5 µM of a 21 bp dsDNA from the Widom 601 sequence for nucleosome assembly known to directly interact with BRG1 (*5*) (right), as compared to dsDNA only as control (left), confirms binding between BRG1_C_ and the 21 bp dsDNA. **(B)** *In vitro* droplet assay using confocal microscopy on mEmerald-10xHis-BRG1_C_ either in the absence (top row) or presence (bottom row) of 5% PEG-8000 (w/v), with the addition of either increasing concentrations of the 21 bp dsDNA from the Widom 601 sequence or a 4 kb dsDNA PCR product of random sequence. The presence of even a small amount of Widom 601 sequence significantly enhances BRG1_C_ droplet formation *in vitro*. **(C)** Histograms of the average droplet area and number of droplets per field of view (FOV) for each condition assayed in the bottom row of (**B**). Scale bar for **B** and **C**: 50 µm. *n* = 10 fields of view for each condition in **C**.

**Figure S12.**
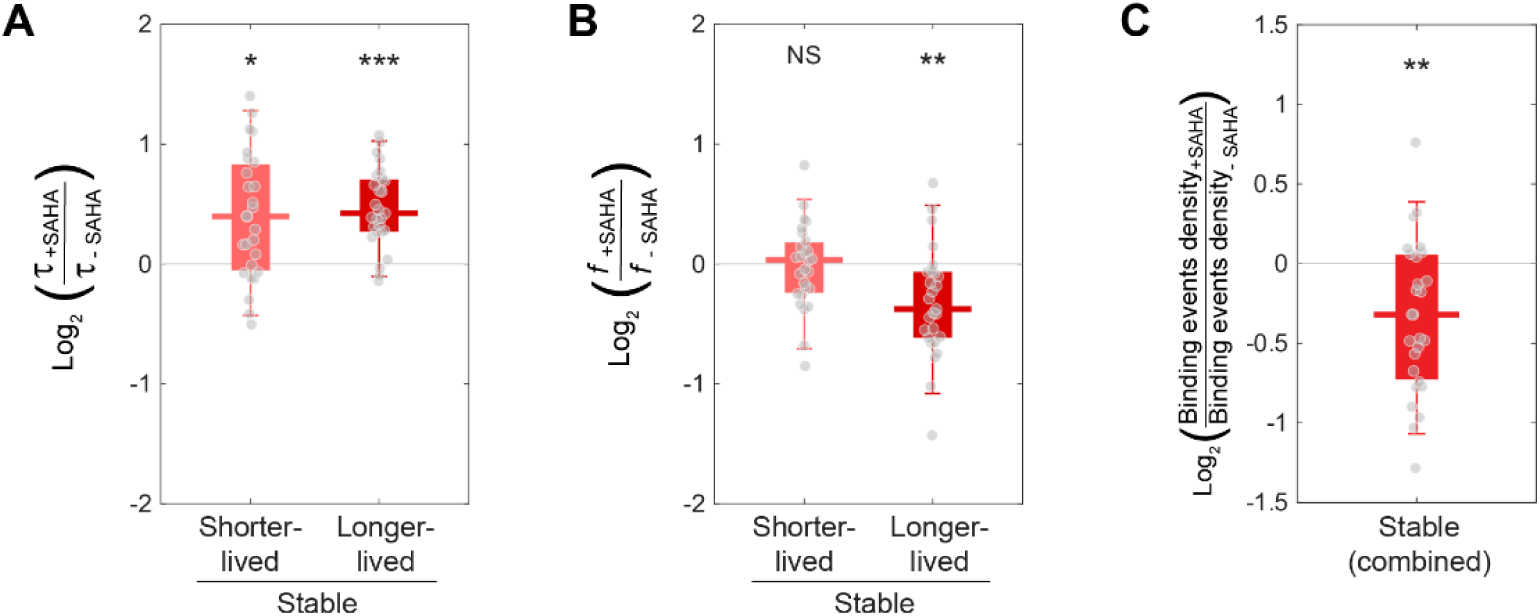
SAHA treatment ascertains that the intranuclear binding dynamics we observed for BRG1_C_ indeed corresponds to chromatin-binding. **(A**–**C)** Box-and-whisker plots of fold differences (on a log_2_ scale), upon SAHA treatment, in binding residence time (τ) **(A)**, corresponding molecular fraction (*f*) **(B)** and binding events density **(C)** for stable chromatin-binding of BRG1_C_, in line with the known role of the bromodomain harbored within BRG1_C_ in contributing to binding hyperacetylated and decondensed chromatin. Each dot denotes a single cell. *n* = 27 and 21 cells for with and without SAHA treatment, respectively.

**Figure S13.**
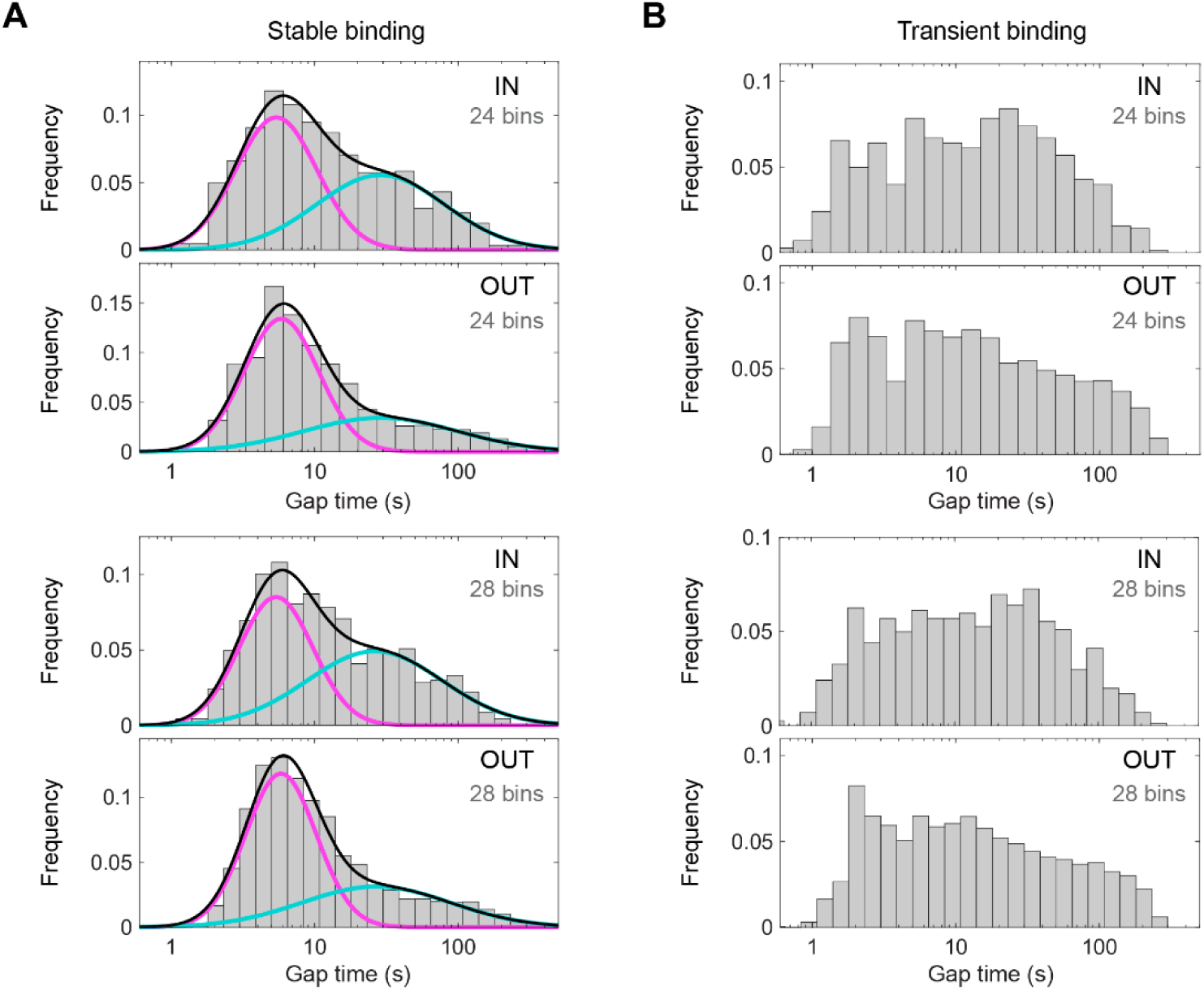
Gap time distribution between transient binding events of BRG1_Cz_ does not exhibit a clear bimodal distribution either inside or outside the condensates. **(A**–**B)** Histograms of gap time between adjacent binding events of BRG1_C_ within a hotspot for stable mode **(A)** and transient mode **(B)**, either inside or outside the nucleolar condensates. In contrast to the gap time between stable binding events which exhibits two temporally distinct peaks that remain invariant upon changing bin size (with the 24-bin case being the one shown in Fig. 5J), the gap time between transient binding events is more homogeneously distributed and cannot be robustly fitted with a bimodal distribution with clearly separated peaks temporally. *n* = 11,660 gap times for **A** and 5,725 gap times for **B**, both from 53 cells.

**Table S1.**
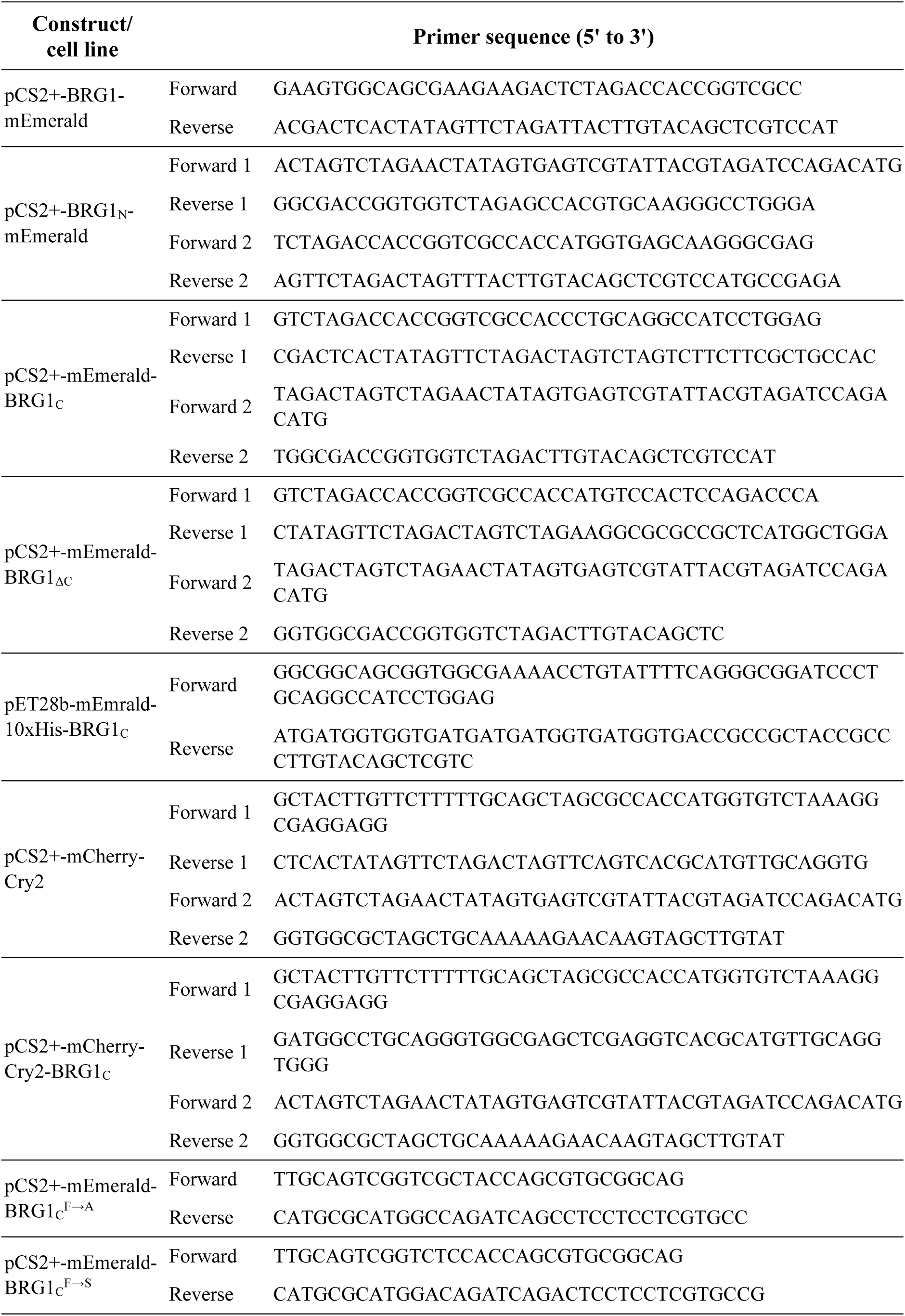
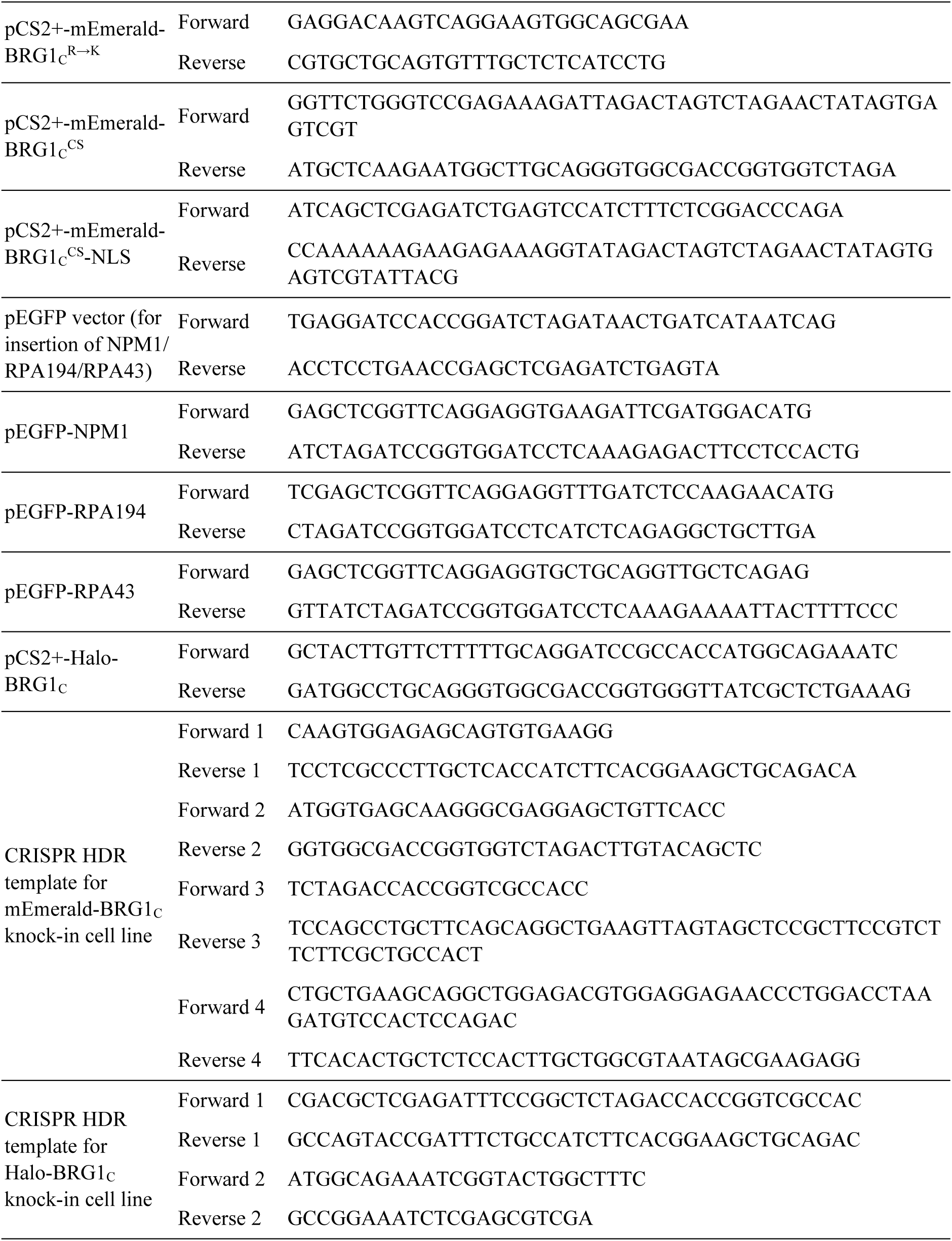
List of primers used for generating constructs and cell lines used in this study.

## Notes

### Competing Interest Statement

The authors have declared no competing interest.

